# Axial patterning of gastruloids via diverse cell type compositions

**DOI:** 10.1101/2025.03.26.645494

**Authors:** Kerim Anlas, Fumio Nakaki, Nicola Gritti, Jordi Font-Reverter, Krisztina Arató, Martí Planasdemunt-Hospital, David Oriola, James Sharpe, Vikas Trivedi

## Abstract

The formation of the germ layers and antero-posterior (AP) axial patterning are interlinked milestones of embryogenesis. Gastruloids, *in vitro* models from aggregated embryonic stem cells (ESCs), permit the study and deconstruction of these events. Gastruloids are successfully generated from ESCs of variable pluripotent states, but it remains unknown how the initial conditions influence cell type composition and to what degree resulting variations in spatial patterning can converge onto an elongated body axis. To address this, we comparatively study aggregates from varying proportions of primed and naive ESCs. Despite differences in AP symmetry-breaking dynamics and distinct trajectories toward either anterior mesodermal or neuro-ectodermal fates, all conditions produce elongated gastruloids. Furthermore, timed modulation of Activin/Nodal signaling in aggregates equalizes AP polarization modes and mixing of initial ESC pluripotent states generates gastruloids with enhanced tissue type diversity. This work therefore uncovers a previously unappreciated developmental flexibility underlying mammalian AP axial patterning.

## Introduction

The bilaterian body plan is established early during embryogenesis as part of a phase termed gastrulation (***1***). Hallmarks of this process are the specification of the three germ layers, ecto-, meso- and endoderm, from an initially pluripotent embryonic stem cell (ESC) population and their spatial organization along (at least) an antero-posterior (AP) body axis (***2,3***). Notably, the emergence and study of *in vitro* embryo-like structures generated from pluripotent stem cells (PSCs) has greatly enriched our understanding of such early developmental events. In particular, aggregates of ESCs, termed 3D gastruloids (***4***), have not only been shown to recapitulate AP symmetry breaking, germ layer specification and axial patterning (***5***), but also to provide insights into the differences in signaling requirements between species at these early, inaccessible stages of embryogenesis (***6***). Consequently, gastruloids are being probed with an ever-increasing array of imaging (***7*,*8***), biophysical (***9*,*10***) and molecular tools including transcriptomic, proteomic and epigenomic technologies (***11–17***).

The current understanding of the spatiotemporal coordination between emerging cell types during mammalian axial patterning is primarily based on observations from the controlled environment of the embryo, where localized signals from extra-embryonic tissues guide ESC differentiation and overall morphogenesis (reviewed in ***18*,*19***). Conversely, gastruloids develop in absence of such tissues and associated cues, hence revealing the inherent “self-organizing” potential of ESCs to generate a (transcriptionally) patterned mammalian body plan. Here, emergence and subsequent polarization of the conserved mesodermal marker Brachyury or T (T) demarcates initial germ layer and AP axis establishment or symmetry breaking, respectively (***7,14,20***). While gastruloids mimic spatial gene expression mostly faithfully, especially along the AP axis, they lack much of the complex morphogenesis and morphology as well as certain tissue (sub-)types of the native embryo. Hence such aggregates arguably recapitulate subsets of the total *in vivo* developmental trajectories, referring to cell and tissue behaviors as well as gene expression and cell state transition dynamics (***18***). This also raises the possibility that, due to the lack of embryo-like mechanical and geometrical constraints, gastruloids can be employed to study a more unconstrained developmental trajectory space of body axis formation (***18*,*21***). Recent research indeed suggests that such trajectories exhibited by gastruloids or, broadly, by ESCs removed from their default context, can occasionally be more flexible than those in the native embryo (***22***), for instance in the case of different transcriptomic ESC states *in vivo* and *in vitro* which nevertheless converge on similar cell fates (***7***). Furthermore, while observations in gastruloids are reproducible, especially within studies, phenotypically similar aggregates can have distinct tissue compositions across labs, as illustrated by varied observations regarding T polarization dynamics and opposing anterior domain identity, i.e. the level of overlap between a Sox2^+^ (pluripotent or, at this stage, likely neurally primed) domain and the emerging posterior T pole during AP polarization (***7,10,14***). Another study reports differentiation biases in individual gastruloids toward either mesodermal or neuroectodermal fates (***15***).

In this context, it should be mentioned that a challenge related to the work with PSCs in general lies in the diversity of adoptable pluripotent states (***23*,*24***), comprising a spectrum of transcriptional (***25*,*26***), epigenetic (***27–29***) and metabolic configurations (***29*,*30***) that bias their response to differentiation cues and thus lineage specification (***28*,*32–36***). For mouse ESCs (mESCs), the two most widely employed types of media for sustained self-renewal are ES-LIF-based (containing animal serum and Leukemia Inhibitory Factor (LIF)) and 2i-based (with small molecule inhibitors targeting GSK-3β and MEK or Src kinase) (***37,38***). While, generally, these two conditions promote distinct compositions of pluripotent cells, where ES-LIF is known to permit heterogeneous pluripotency levels and 2i facilitates a more uniform naive population (***26***), the constituent base reagents may also differ across labs (***7***). The effects of the initial mESC (pluripotent) state on gastruloid morphology and tissue composition are just beginning to be scrutinized (***39,40***), in particular, it remains unclear how AP polarization and spatial germ layer patterning dynamics are impacted. Notably, when considered jointly with the above mentioned variation in aggregate tissue composition, this presents a unique opportunity to uncover the variety of emerging combinations of cell types that are able to form an elongated AP axis - beyond merely identifying the sources of variance encountered across labs. We remark that such variability in gastruloid developmental trajectories does not (necessarily) refer to distinct molecular mechanisms, but rather points to what subsets of embryonic trajectories are, in principle, sufficient to generate an axially patterned body plan (***18***). Experimentally addressing this by leveraging differentiation biases imparted by the aggregated cell population can therefore elucidate minimal principles underlying the generation of animal form *in vivo* and potentially identify new strategies to engineer desired cell types *in vitro*.

## Results

In order to comparatively study gastruloid AP polarization, elongation and germ layer patterning upon variation of source mESC pluripotent state composition, we initially focused on T expression dynamics as a useful proxy for these processes (***4*,*5*,*7*,*12*,*14,20***). Hence, we generated an endogenous T-reporter double-transgenic (2KI) G4 mESC line, additionally featuring ubiquitous nuclear fluorescence (T-p2a- H2B-eGFP and CAG::H2B-mKO2, **Fig. S1A**, also see Materials & Methods). For elucidating the influence of initial PSC state on biasing lineages under equivalent differentiation protocols (with N2B27 medium and external Wnt upregulation via a CHIR99 pulse over 24 hours (h)), we cultured 2KI mESCs in three distinct conditions prior to aggregation **(Fig. 1; Fig. S1B,C)**: (i) “base medium” which permits the presence of primed pluripotent subpopulations, facilitating heterogeneous pluripotency levels; (ii) base medium with the MEK/ERK inhibitor PD03 to decrease primed cell admixture (base +1i); and (iii) base medium with Src-kinase inhibitor CGP77 and GSK-3B inhibitor CHIR99 (base+2i) to uniformly confine cultured mESCs to a homogeneous, naive pluripotent state (***26*,*41;*** for detailed formulations see Materials & Methods). We here opted for CGP77 (termed “alternative 2i”) instead of the more commonly employed PD03, given that traditional 2i medium has been shown to incur irreversible epigenetic and genomic alterations in ESCs, leading to diminished developmental potential (***42,43***). For gastruloids from base+2i, the Wnt agonist pulse had to be delayed by 24h to ensure AP polarization and elongation, that is, from 48-72h post-aggregation (hpa) to 72-96hpa. **(Fig. S1D)**.

**Fig. 1.**
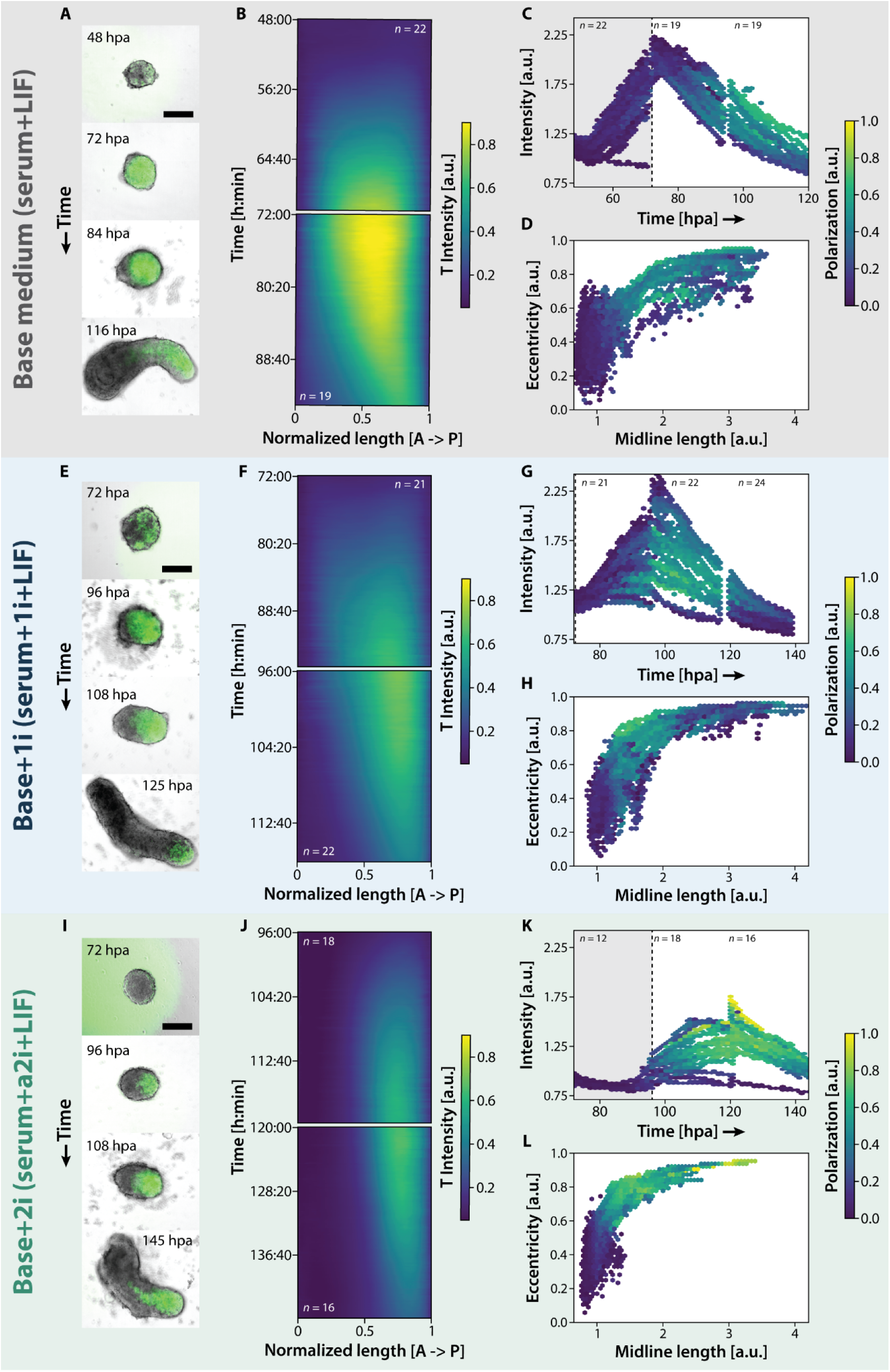
Initial medium condition influences T polarization and aggregate elongation dynamics. **A), E), I)** Representative images of gastruloids from mESCs cultured in either base medium (serum + LIF), base+1i (serum+LIF+0.3μM PD03) or base+2i (serum+LIF+3μM CHIR99+1.5μM CPG77), facilitating distinct pluripotent states spanning heterogeneous to homogeneous, naive pluripotency. The images depict the onset of T expression, polarization and maximum elongation for each condition. Scale bars = 200μm. **B), F), J)** Kymographs displaying average T expression intensity over time along the normalized major or AP axis of the gastruloids (as denoted by T expression) from each medium condition (or pluripotent state). **C), G), K)** Hexbin (2D histogram) plots of individual gastruloid T intensity over developmental time for each condition. The color code indicates the degree of (T) polarization. The shaded area indicates the time window of CHIR99 application (in **G**, CHIR99 was applied prior to the displayed time frame). **D), H), L)** Hexbin plots of individual aggregate eccentricity (describing elongation) over midline length (along AP axis) for each condition.

### Initial medium condition influences T polarization and aggregate elongation dynamics

Wide-field imaging of developing aggregates reveals clear differences in patterns and dynamics of T emergence between the three media conditions **(Fig. 1; Figs. S1 & S2)**: In base **(Fig. 1A,B)** and, to a lesser degree, base+1i **(Fig. 1E,F)**, the reporter initially displays a ubiquitous expression which then polarizes to define the AP axis. In contrast, in base+2i, T emerges with a notable bias to one side of the aggregate, immediately denoting the AP axis **(Fig. 1I,J).** Interestingly, the increase in T intensity following the CHIR99 supplementation is also distinct across conditions **(Fig. 1C,G,K)**, with base+1i and base+2i exhibiting more heterogeneous and diminished profiles, respectively, suggesting a disparate response to differentiation cues depending on starting conditions or state. Moreover, in gastruloids from base medium, high T polarization (>0.5, see Materials & Methods) typically occurs only after overall T intensity decreases. Conversely, in base+2i, this degree of polarization can be reached before or at the same time as maximum T expression. Such distinctions further highlight how the inferences of cellular interactions and fate decisions from *in vitro* models across labs require careful scrutiny as patterning dynamics and external signal requirements are remarkably dependent on initial PSC state.

Furthermore, different conditions have variable dependencies on CHIR99 treatment for T expression and symmetry-breaking (base: 54/55 across 2 batches; base+1i: 13/59 across 3 batches; base+2i: 39/60 across 3 batches). **(Fig. S1E; Fig. S2A,B,E,F,I,J)**. Gastruloids from base medium also display larger batch-to-batch variation in the timing of T polarization: ⅗ batches polarize by 48 hours post-aggregation (hpa) as opposed to ⅖ by 72hpa **(Fig. S1F,G)**. Such differences in gene expression are also paralleled in gastruloid rheological properties at 24hpa based on fusion kinetics (**Fig. S1H, *44,45***). Compared to base+1i and 2i, mESCs from base medium generate aggregates that undergo complete fusion, implying a more fluid-like behavior. Thus, gastruloids from different conditions are in different mechanical states prior to CHIR99 application. These distinctions in tissue-level rheology emanate from differential cell-level properties including adhesion, migration and neighbor exchange (***44***) which are likely to contribute to subsequent variations in aggregate patterning dynamics. While gastruloids across conditions can undergo axial elongation (quantified by eccentricity and midline length, see Materials & Methods) with CHIR99 treatment, the extent thereof varies significantly **(Fig. 1D,H,L)**: Aggregates from base+1i more consistently reach the most elongated phenotypes while base medium-derived samples exhibit more variability. Interestingly, for base and base+1 media, changes in the strength of T polarization are not concomitant with changes in gastruloid eccentricity, i.e. shape change does not precede AP polarization, unlike for base+2i. It is conceivable that such observations are linked to the gastruloids featuring distinct cellular compositions between conditions.

Having identified distinctions in T polarization dynamics on a tissue-level, we then set out to explore possible differences on a single-cell level. To that end, we employed light sheet fluorescence microscopy (LSFM) on sparsely labeled gastruloids from each source mESC state during AP polarization (**Fig. S3A; Movies S1-3,** see Materials & Methods). This was followed by automated 3D tracking to elucidate T state transitions and migratory dynamics. Individual cells were thus categorized as either T^+^, T^-^ or T^mid^ (an intermediate state, see Materials & Methods). The average mean squared displacement (MSD) exponent of cells shows that throughout all datasets and T states, cells are in diffusive to slightly superdiffusive motion during symmetry breaking **(Fig. S3B)**. Moreover, T^+^ cells have slightly higher MSD exponents than T^-^ and T^mid^ populations on average which is consistent with increased motility in T^+^ states **(Fig. S3C)**. We then looked into cell state transitions during AP polarization and found that all conditions feature a significant proportion of cells that show fluctuating T expression **(Fig. S3D-G)**. Focusing on more stable transitions, in aggregates from base medium, many T^+^ cells convert to T^-^ ones which often migrate toward the anterior (along with pure T^-^ cells), unlike the mostly posterior but (more) random movement of T^+^ cells **(Fig. S3E)**. This directional migration was not clear in other conditions. Gastruloids from base+1i and particularly base+2i medium had fewer pure T^+^ cells and less state transitions, although base+1i displayed slightly more T^-^ to T^+^ conversions than base medium **(Fig. S3F-G)**.

### Tissue fate specification and localization differs between initial media conditions

Prompted by the above findings, we hypothesized that the observed distinctions in T AP polarization and elongation modes may be based on different proportions of progenitor populations being generated concomitantly. To examine this, we comparatively performed *in situ* HCR of key germ layer and axial patterning markers in aggregates from wild-type (WT) G4 mESCs **(Fig. 2; Fig. S4)**. Gastruloids from base medium can display a polarized *T* domain and expression of the definitive endoderm marker *Sox17* already by 48hpa, i.e. before CHIR99 application, consistent with previous reports (***7***) **(Fig. 2A)**. At this stage, *Aldh1a2*, a marker for lateral plate mesoderm, exhibits weak expression, and *Eomes*, denoting anterior meso- and endodermal lineages, overlaps with T **(Fig. 2A)**. Hence differentiation into the earliest mesodermal subtypes and the initial separation of anterior (Eomes^+^) versus posterior (T^+^) mesoderm is not completed by 48hpa. At 72hpa, two distinct domains - *Aldh1a2*^+^ and *Eomes*^+^ - segregate anteriorly from the posterior T pole **(Fig. 2B)**. *Sox17* is now more broadly expressed, while *Sox2*^+^ cells, which are either undifferentiated or neuroectodermally-primed, can be identified throughout the aggregate **(Fig. 2B)**. The 72hpa time point also marks the onset of localized *Gata6* expression at the anterior end of the aggregate **(Fig. S4A)**, partly overlapping with the *Eomes* domain. *Gata6* is one of the earliest markers for (pre-)cardiac mesoderm and endoderm (***46***), arguing that cardiovascular tissue is starting to be specified in this domain. Between 96-120hpa (batch-dependent), when base medium-derived gastruloids reach maximum elongation, *Gata6* expression further intensifies **(Fig. S4B)**. Furthermore, neural progenitor cells (NPCs) can be identified by expression of *Sox1* together with *Sox2* **(Fig. 2C)** which are distributed along the AP axis of the gastruloid, though not always in the anteriormost region, which features domains of mostly *Gata6*^+^ and *Eomes*^+^ but also *Sox17*^+^ cells (20/22 over two HCR batches) **(Fig. S4B).** Additionally, the somitic mesodermal marker *Meox1* (***47***) is broadly located in the trunk region of the gastruloid, adjacent to *Sox1*- and *Sox2*- expressing regions **(Fig. 2C)**.

**Fig. 2.**
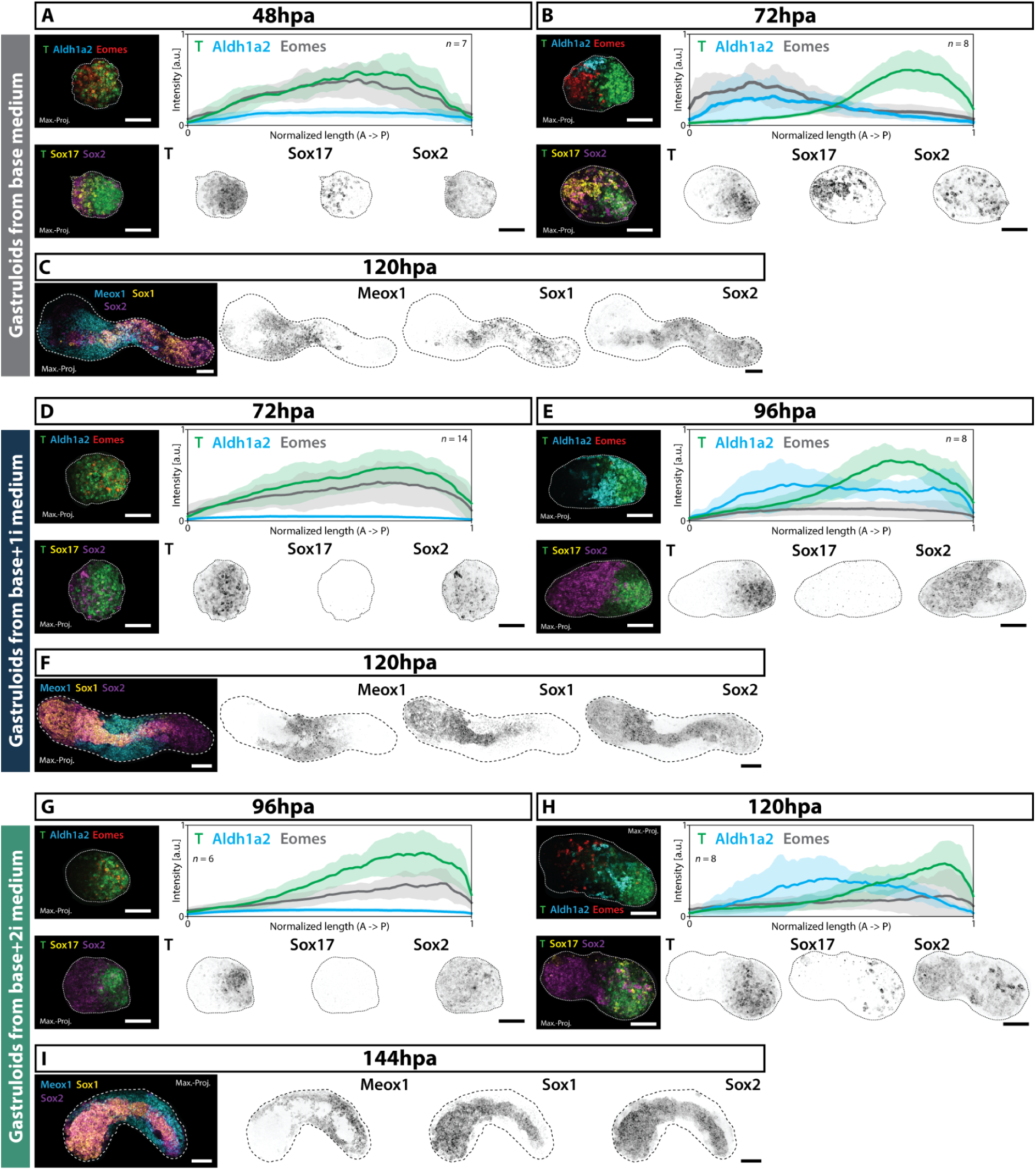
Tissue fate specification and localization differs between initial media conditions. Representative images (maximum intensity projections) of HCR stainings of germ layer and axial patterning marker genes in gastruloids from three mESC media conditions (or pluripotent states) at comparable developmental timepoints: Early T or AP polarization (48hpa in base medium - **A)**, 72hpa in base+1i - **D)**, 96hpa in base+2i - **G)**); late polarization or early elongation (72hpa in base medium - **B)**, 96hpa in base+1i - **E)**, 120hpa in base+2i - **H)**); and maximum elongation (120hpa in base medium - **C)**, 120hpa in base+1i - **F)**, 144hpa in base+2i - **I)**). All scale bars = 100μm. Line plots in top right panels of **A), B), D), E), G), H)** depict the normalized fluorescence intensity of marker genes T, Aldh1a2 and Eomes (based on sum intensity projections) along the aggregate’s normalized major or AP axis (as denoted by T expression). Central lines denote the average intensity, the surrounding shading marks the standard deviation across replicates.

In contrast to aggregates from base medium, gastruloids from base+1i and 2i mostly lack anterior mesodermal lineages, instead containing NPC populations in the anterior region. Such a remarkable difference in spatial patterning manifests early on in development. In both conditions, mesendoderm specification does not occur prior to CHIR99 treatment as demonstrated by lack of *T* and *Sox17* signal at 48hpa for base+1i **(Fig. S4D)** and 72hpa for base+2i **(Fig. S4E)**. In gastruloids from base+1i, *Eomes* expression is significantly lower at 96hpa when T is clearly polarized, unlike the case in base media **(Fig. 2D,E).** *Aldh1a2*, though strongly expressed, does not localize to the anteriormost part **(Fig. 2E)**, and *Sox17* transcripts are extremely sparse, if not absent **(Fig. 2D,E; Fig. S4C)**. In contrast, *Sox2* is broadly expressed at 72 and 96hpa, both overlapping with the T domain and also exclusively present in the anterior T^-^ half of the polarized gastruloid. Due to the lack of co-expression with any of the assayed markers in this region, it likely contains neuroectodermally-primed cells. Indeed, at maximal gastruloid elongation (reached around 120hpa), the anteriormost part is mostly *Sox1*+ and *Sox2*+ double positive, denoting an abundance of NPCs **(Fig. 2F)**. Conversely, anterior *Gata6* expression could not be observed (0/11 aggregates, **Fig. S4C**), whereas a broad domain of *Meox1*+ somitic mesoderm is found across the gastruloid trunk **(Fig. 2F)**.

In gastruloids from base+2i, similar to those from base+1i and unlike in base medium, *Aldh1a2* is expressed in the trunk, but not the anterior region by 120hpa, ie. 48h after initial application of CHIR99 **(Fig. 2H)**. However, both *Eomes* and *Sox17* expression is higher by this stage than in base+1i **(Fig. 2E,G,H)** yet not localized to the anterior end. A broad *Sox2*^+^ domain with strong *Sox1* expression overlap forms the anteriormost domain upon reaching maximum elongation by 144hpa **(Fig. 2I)** indicating enrichment of NPCs. Anterior *Gata6* signal could only be identified in a few aggregates (4/19) **(Fig. S4F)**. As in all conditions, *Meox1*^+^ somitic mesoderm is found throughout the gastruloid trunk, adjacent to the NPCs **(Fig. 2I)**. These distinct spatial arrangements of gene expression domains show a previously unappreciated diversity in the axial patterning of gastruloids resulting from lineage biases due to alterations in starting culture conditions.

### Differential generation of anterior mesodermal versus neuroectodermal populations

Prompted by the above findings, we sought a more detailed comparison of the developing cell lineages between the three initial states via single-cell RNA sequencing (scRNAseq) on the source mESCs and on gastruloids pre- and post-CHIR99 treatment as well as at maximum elongation, i.e. 72h after initial CHIR99 supplementation **(Fig. 3; Fig. S5)**. Uniform Manifold Approximation and Projection (UMAP) of our larger, time-resolved gastruloid dataset yields 16 distinct cell populations, spanning naive to primed pluripotent states as well as derivatives of all germ layers **(Fig. 3A)**. In general, labeling cells by time points **(Fig. 3B)** illustrates a greater degree of visual separation in UMAP space than labeling by media condition **(Fig. 3C)**. We identified a naive pluripotent cluster (Cluster #1) and four primed pluripotent populations (#2-5) with subtle differences in expression levels and prevalence of markers like *Cd63, Utf1, Pim2, Dnmt3b*, and adhesion gene *Cdh1* **(Fig. 3A,D)**. A more detailed and separate UMAP analysis of mESC populations from the three conditions corroborates that base medium facilitates a heterogeneous mix of pluripotent states **(Fig. S5A-D)**, including both naive and primed populations, consistent with previous findings (***25*,*26***). The various clusters which are situated across the pluripotency spectrum are differentially dominated by mESCs from the three employed media: for instance, one naive pluripotent cluster (II, expressing *Dppa5a*, *Klf4* and metabolic genes *Gpi1*, *Pgk1* and *Aldoa* which encode glycolytic enzymes) is almost entirely derived from base+2i (∼99.5%) that exclusively promotes naive pluripotent states in general, whereas base medium contributes predominantly (∼88%) to a primed pluripotent cluster (V, expressing *Pim2, Dnmt3b, Pou3f1, Lefty1* as well as epithelial markers *Krt8* and *Krt18).* The base medium condition indeed features more cells with higher levels of differentiation markers, including early mesodermal (*T, Fgf5, Wnt3*) and neuroectodermal (*Sox1, Sox3*) genes **(Fig. S5A)**. Base+1i-grown mESCs show mostly naive states (high levels of *Pou5f1/Oct4*, *Nanog* and *Sox2*) with limited admixture of primed subpopulations.

**Fig. 3.**
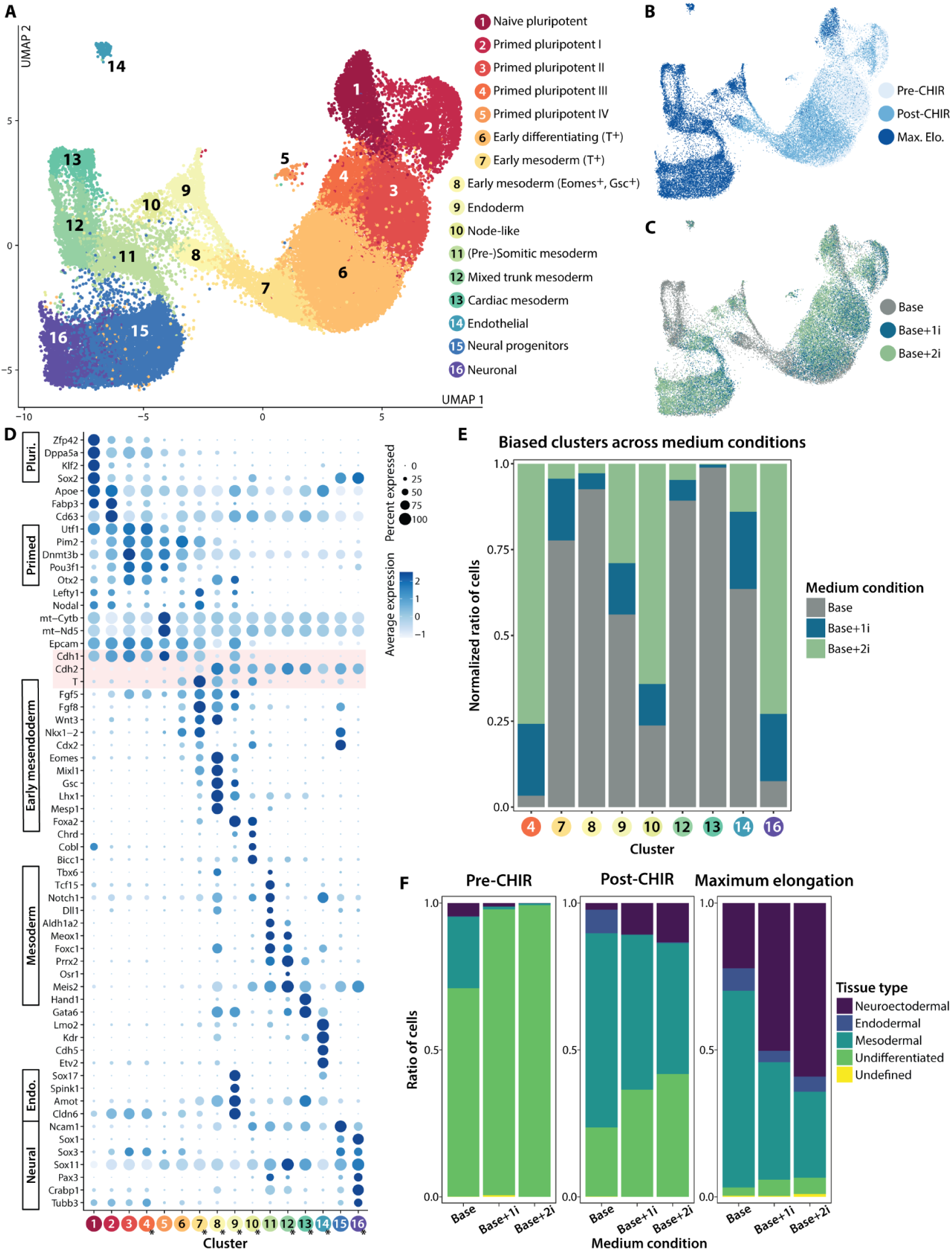
Single-cell transcriptomics delineate distinctions in lineage specification dynamics. **A)** UMAP plot of the integrated gastruloid dataset spanning three initial mESC media conditions at three comparable developmental stages. Color code is according to cluster analysis (see labels on the right). **B)** UMAP with each cell color coded according to the developmental stage. **C)** UMAP color coded according to the medium condition. **D)** Dotplot displaying scaled average expression of numerous marker genes denoting cell state and the percentage of cells expressing a given marker for each cluster in **A)**. Highlighted in red are key cell adhesion and EMT markers Cdh1 and Cdh1 as well as T. Clusters marked with an asterisk represent clusters with over- (or under-) representation of a given initial media condition. **E)** Bar chart of all “biased” clusters from **A)** depicting the normalized ratio of cells from each mESC media condition. **F)** Bar plots showing the ratio of cells from each germ layer or tissue type for each individual dataset (i.e. from a given medium condition) at each developmental stage.

In developing aggregates, cells exiting pluripotency cluster into the “early differentiated” population (#6), expressing early mesodermal markers *T, Fgf8, Wnt3, Nkx1-2* **(Fig. 3D)**. Fully differentiated mesodermal cells (#7) display higher levels of these markers plus *Eomes*, *Cdx2*, and *Mixl1*. An early anterior mesoderm cluster (#8) exhibits elevated *Wnt3, Eomes, Gsc, Lhx1, Mesp1,* and lower *T* and *Fgf8*. The “T-high” mesoderm cluster (#7) contrasts with the “Eomes-high” mesoderm (#8) in terms of cell adhesion marker expression profiles: T-high has elevated *EpCAM* and *Cdh1* but low *Cdh2*, while Eomes-high shows low *Cdh1*, low *EpCAM*, and increased *Cdh2*. These adhesion differences may drive the segregation of the posterior T pole from the anterior Eomes domain in base media-derived gastruloids, consistent with recent reports (***8*,*48*,*49***). We also find a prominent endodermal group (#9, marked by *Foxa2, Sox17, Spink1, Cldn6*) and a small node-like cluster (#10, high in *Foxa2, Chrd, Cobl, Bicc1*). There are three additional mesodermal populations: #11 (pre-)somitic mesoderm (*Aldh1a2, Meox1, Notch1*); #12 mixed trunk mesoderm (*Meis2, Sox11, Osr1*); and #13 cardiac mesoderm (*Gata6, Hand1*). Endothelial progenitor cells (#14, *Lmo2, Kdr, Cdh5, Etv2*) are also present. Of the two neuroectodermal clusters, one (#15) includes NPCs (marked by *Ncam1, Nkx1-2, Sox1, Sox2* and *Sox3*) and the other (#16) likely contains more committed neuronal progenitors, exhibiting stronger and more pervasive (>75%) expression of *Sox1* as well as markers *Pax3, Crabp1* and *Tubb3*.

Together these populations represent the diverse cellular states that emerge during gastruloid development across the three conditions. To identify key distinctions, we focused on those 9 of the 16 clusters that showed significant ‘bias’, with one medium contributing either most of the respective cells (>50%) or very few (<10%). The early mesodermal clusters (#7,8) appear to be dominated (>75%) by cells from base medium-derived gastruloids, featuring less contribution from base+1i and base+2i **(Fig. 3E).** Particularly the (early) anterior mesoderm (#8) is strongly biased (∼93%) towards base medium, consistent with the HCR stainings **(Fig. 2)**. The endodermal population (#9) is also enriched for base-derived cells (∼56%), although distinctions are not as drastic as for the previous clusters **(Fig. 3E)**. Similarly, trunk (#12, ∼89%), and cardiac (#13, ∼99%) mesodermal subtypes are almost exclusive to base medium, corroborating observations from the *Gata6* stainings **(Fig. S4).** While the endothelial cluster (#14) is moderately enriched in base medium (∼64%), node-like (#10, ∼64%) and neuronal progenitor (#16, ∼73%) cells are dominated by base+2i-derived gastruloids. These results highlight how medium composition shapes the landscape of cellular interactions and response to signals that bias germ layer proportions and cell-type distributions across conditions.

To further elucidate germ layer specification dynamics, we analyzed the proportions of ectodermal, mesodermal, and endodermal cells for each source mESC state at the assayed developmental stages **(Fig. 3F, Fig. S5E)**. Before Wnt stimulation, most cells are undifferentiated, with only base medium-derived gastruloids featuring significant mesoderm (∼24% vs. <1% in other media; **Fig. 3F**). After CHIR99 treatment, there is a considerable increase in mesodermal cells in aggregates across conditions (∼66% in base, ∼53% in base+1i, and ∼45% in base+2i). Neuroectoderm is present in all conditions, but is highest in base+1i (∼11%) and base+2i (∼13%). At this stage, gastruloids from base+2i also contain the most undifferentiated cells (∼42%). Endoderm is most notable in base medium (∼8%). By maximum elongation, mesoderm remains dominant in the base condition (∼67%), while neuroectoderm is highest in the other two (>50%). Interestingly, the endodermal populations have moderated across conditions (∼8% in base medium, ∼4–5% in others), as also reflected in the Sox17 stainings **(Fig. S4C,F)**. Altogether, there is a remarkable difference in the cellular compositions between the initial conditions: Base medium-derived gastruloids develop predominantly mesodermal (and endodermal) tissues, whereas aggregates from base+2i have comparatively more neuroectodermal tissues, despite the external Wnt activation by CHIR99. In spite of this variability in cell type compositions, Hox gene expression across media conditions at maximum gastruloid elongation is qualitatively similar **(Fig. S5F)**, further illustrating how a transcriptionally patterned, elongated AP axis is nevertheless successfully generated, independent of starting ESC state.

### Upregulation of Activin/Nodal signaling equalizes symmetry breaking dynamics between initial pluripotent states

The observed distinctions in germ layer specification dynamics imply that the gastruloids from the three starting mESC states explore different regions of the differentiation landscape since their constituent cells exhibit different responses to signals. Therefore, we asked if we can shift tissue proportions and the mode of symmetry-breaking in gastruloids from one condition to mimic those in another through external signals. We focused on the formation of an anterior *Eomes* domain that is reproducibly observed only in base medium-derived gastruloids and highlights a key distinction in AP patterning between conditions. Unlike the gastruloids from base+1i- and base+2i-grown mESCs, those from base medium spontaneously develop *Eomes* domains even without CHIR99 (43/51, **Fig. S6)** suggesting an innate regulation of the levels of relevant signals to promote anterior mesendoderm fate. Studies have demonstrated that the pluripotent epiblast under Nodal/Smad2/3 signaling induces *Eomes*, leading to the specification of the anterior mesoderm and definitive endoderm lineages, also via restriction of T activity during early gastrulation (***50,51***).

Average expression in our data shows that in both base+1i- and base+2i-derived gastruloids, Nodal levels are lower than in aggregates from base medium before CHIR99 treatment **(Fig. S5A)**. Thus we added Activin A (ActA, 50ng/ml) alongside the CHIR99 pulse to the former gastruloids which induced a robust anterior *Eomes* domain in nearly all samples (21/22 for base+1i, 19/19 for base+2i), resembling those from base medium with CHIR99 alone (36/39) **(Fig. 4A-D; Fig. S6)**. Without ActA, anterior *Eomes* domains were infrequent (8/32 for base+1i, 11/31 for base+2i), albeit variable across batches **(Fig. 4E-H)**. Strikingly, such changes in fate decisions via ActA treatment impact the dynamics of T polarization as well: Similar to base medium-derived gastruloids, ActA and CHIR99 co-treatment promotes initially ubiquitously distributed T expression, prior to eventual restriction and AP polarization, particularly in base+2i gastruloids **(Fig. 4B,D versus 4F,H)**. These results indicate that reduced Activin/Nodal signaling in aggregates from base+1i and base+2i conditions during mes(- endo)derm induction limits the formation of an anterior *Eomes* domain.

**Fig. 4.**
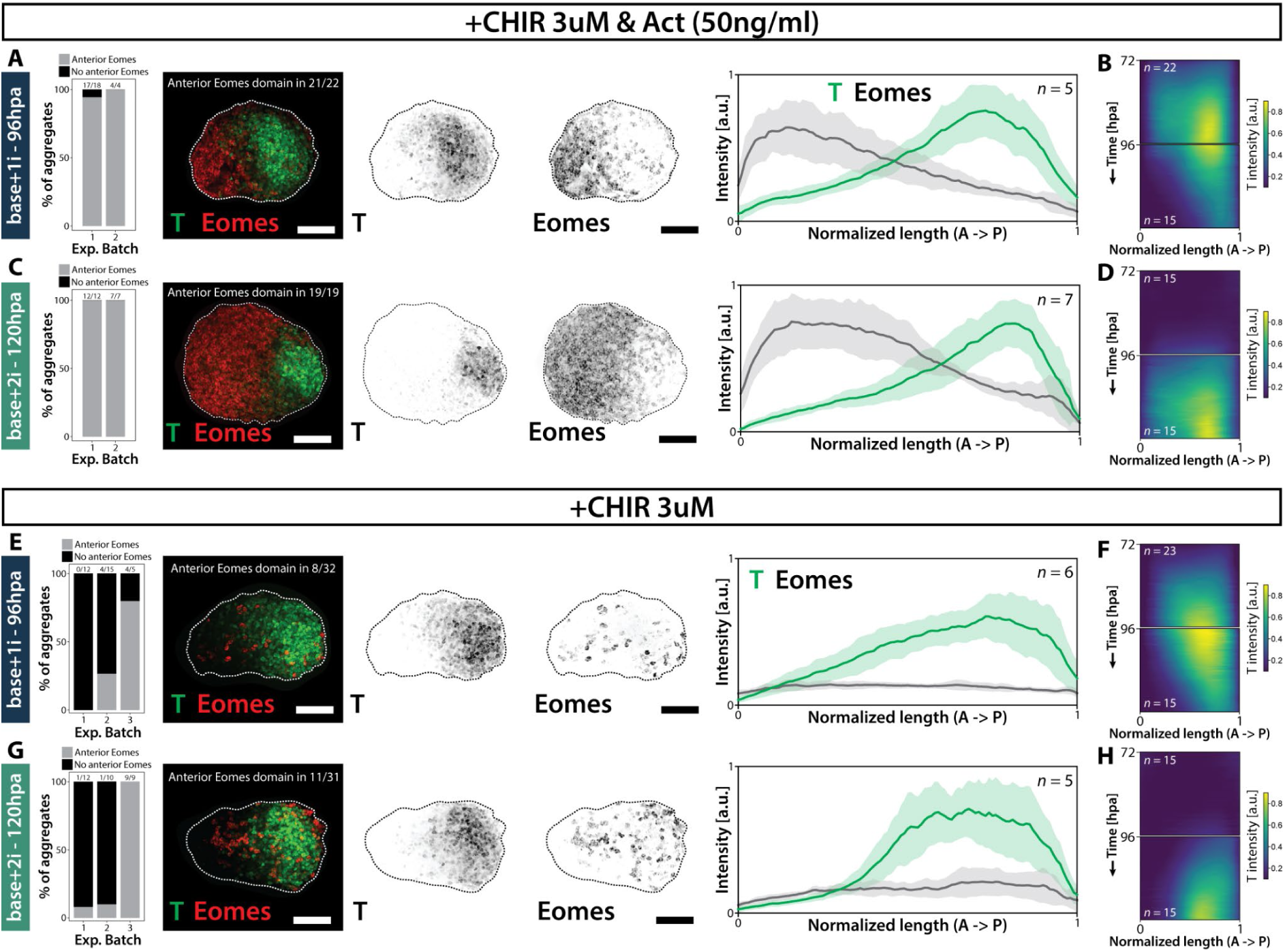
Upregulation of Activin/Nodal signaling equalizes symmetry breaking dynamics between initial pluripotent states. Gastruloids from mESC cultured in either base+1i (**A), B); E), F)**) or base+2i (**C), D); G), H)**) were subjected to a double pulse of CHIR99 (CHIR) and ActivinA (Act, **A) - D)**) instead of the normal, CHIR99-only pulse (**E) - H)**). Leftmost panels in **A), C), E), G)** depict bar charts showing the percentage of aggregates from a given experimental batch exhibiting an anterior Eomes domain. The middle panels show representative images (maximum intensity projections) of HCR stainings of T and Eomes. Scale bars = 100μm. The rightmost panels display line plots of normalized fluorescence intensity of marker genes T and Eomes along the gastruloid’s AP axis (as demarcated by localized expression of T). Central lines denote the average across replicated, surrounding shades the standard deviation. Quantifications are based on sum intensity projections of the HCR data. **B), D), F), H)** Kymographs depicting averaged T expression intensity along the aggregate’s AP axis over developmental time for each condition.

### Mixing of mESCs grown in distinct media yields aggregates with enhanced tissue diversity

Given that non-localized application of a signal (e.g. ActA) can bias the differentiation into certain cell fates (anterior mesoderm) at the expense of other tissues (potentially neuroectoderm), strategies to enhance the diversity of cell fates in “default” gastruloids must rely on minimal external inputs. Furthermore, it is also clear that mESCs from different starting conditions retain their bias towards a lineage in gastruloids despite CHIR99 treatment, unless presented with a new signal (e.g. ActA). Therefore, we asked if we can generate a single gastruloid with cardiac mesoderm, neuroectoderm and endoderm by mixing mESCs from base medium (higher mesodermal diversity) with base+2i mESCs (increased NPC generation) at a 1:1 ratio **(Fig. 5)**. These gastruloids, treated with CHIR99 between 48- 72hpa and analyzed at 120hpa, displayed more frequent anterior *Gata6* expression (18/26 over 2 batches of HCR) as opposed to base+2i- (4/19) or base+1i-derived (0/11) aggregates **(Fig. 5A,B; Fig. S4C).** *Sox2* (mostly NPCs by this stage) signal is now present in anterior regions **(Fig. 5A)**, similar to the base+2i condition, and unlike in base medium where *Sox2* is mostly restricted to the posterior and trunk at maximum elongation **(Fig. 5B)**. *Sox17*^+^ endodermal staining also tends to be more prevalent than in gastruloids from base+2i, but less than in those from base medium (see also **Fig. S4B,F)**. This argues that, in these “mixed” aggregates, cell type proportions are altered to generate the tissues typically present in gastruloids from either of the two source media. Importantly, such a simple mixing can give rise to anterior cardiovascular mesoderm and neuroectoderm more reproducibly than exclusively base+2i or base media-derived gastruloids.

**Fig. 5.**
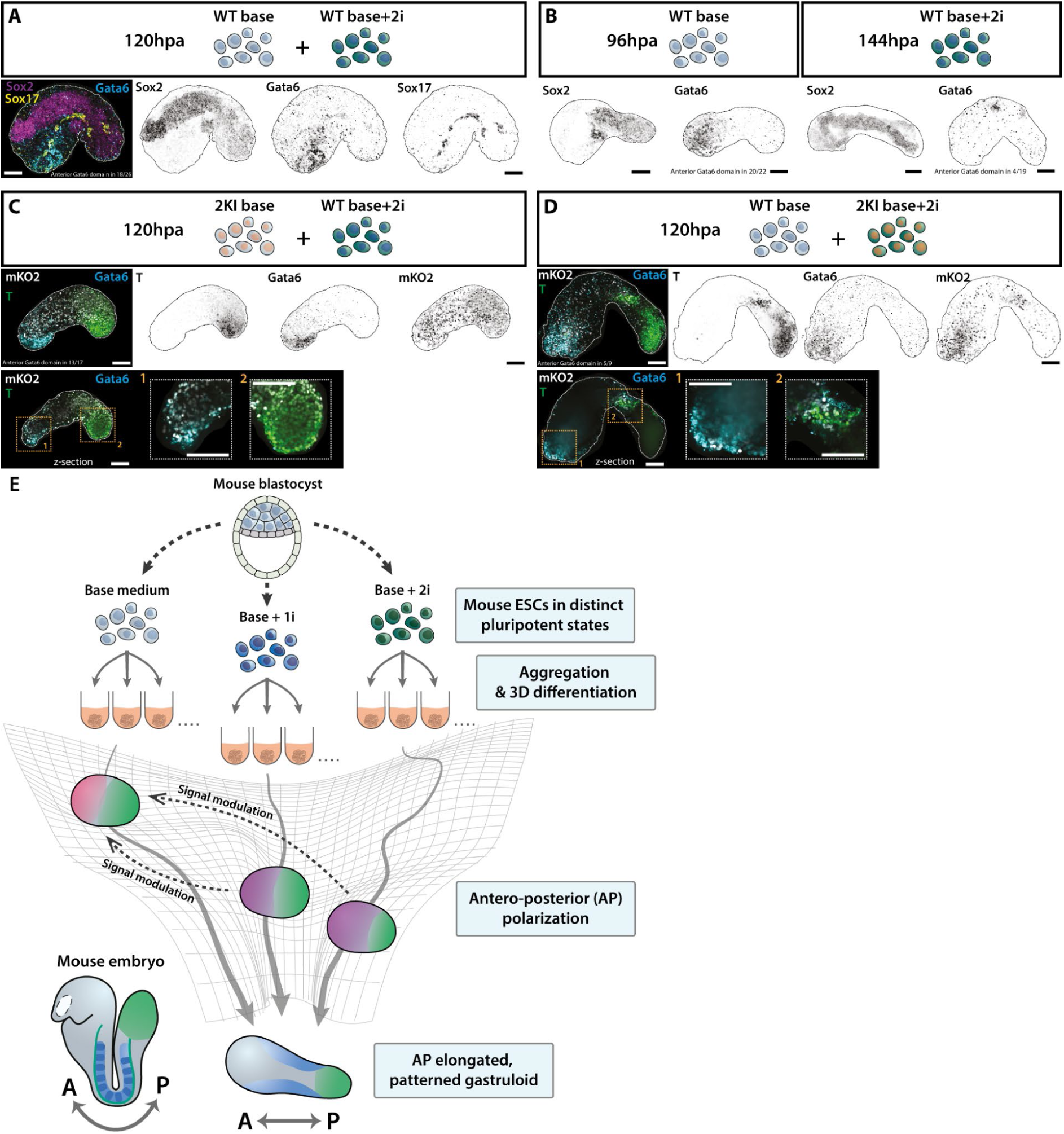
Mixing of mESCs grown in distinct media yields aggregates with enhanced tissue diversity. **A)** Representative images (maximum intensity projections) of HCR stainings of a gastruloid at 120hpa, generated from 1:1 mixed base medium- and base+2i-grown mESCs, showing spatial expression of key germ layer markers. **B)** Representative images (maximum intensity projections) of HCR stainings of gastruloids at 96hpa from base medium (left panels, 2 different specimen) and gastruloids at 144hpa from base+2i (right panels, 2 different specimen), depicting marker genes for comparison with **A)**. **C)** Representative HCR stainings of a gastruloid at 120hpa from WT mESCs from base+2i mixed 1:1 with 2KI double reporter mESCs from base medium. The top panels are maximum intensity projections, the bottom panels are single z-sections to highlight gene expression overlap. mKO2 denotes the fluorescent signal from the ubiquitous nuclear reporter of the double KI line. **D)** Representative images of HCR stainings of a gastruloid at 120hpa from WT mESCs cultured in base medium mixed 1:1 with 2KI reporter mESCs grown in base+2i. Top and bottom panels are structured as in **C)**. All scale bars = 100μm. **E)** Schematic summarizing key findings of this study, highlighting a developmental flexibility in generating a patterned AP axis. Associated trajectories can be manipulated via signal(ing) modulations (amongst others), e.g. via morphogens or cell mixing.

In order to track the contributions of the source mESC populations to the anterior *Gata6* expression domain, we labeled the respective cells through 1:1 mixing of 2KI double reporter with WT mESCs. In both combinations, i.e. base-2KI with 2i-WT **(Fig. 5C)** or base-WT with 2i-2KI **(Fig. 5D)** gastruloids developed *Gata6* expression at the anterior end at 120hpa (13/17 and 5/9 respectively), similar to the base-WT with 2i-WT case. Furthermore, the labeled cells overlap with anterior *Gata6* in both configurations, indicating that base+2i mESCs can contribute to anterior mesodermal fates when mixed with base medium cells. This suggests that cell-cell interactions can alter the signaling environment and thus differentiation bias of source mESC states, similar to the exposure to external signals **(Fig. 4).**

## Discussion

Our findings elucidate the diversity of cell types and their spatiotemporal arrangement in gastruloids generated from different starting mESC populations and thereby provide a framework to unify observations of AP patterning across labs. Given the potential of gastruloid systems to probe, perturb, and engineer events in early mammalian embryogenesis understanding their dependence on initial conditions is essential to establish them as *bona fide* models for both biomedical (***52*,*53***) and tissue-engineering-related applications (***54***). In summary, we show that the mode and dynamics of AP symmetry breaking, marked by T expression, vary depending on initial mESC medium condition and hence pluripotent state. Differences in AP polarization were also linked to distinct elongation behaviors, revealing a previously unreported flexibility of the relationship between AP polarization (molecular symmetry breaking) and shape change (physical-geometric symmetry breaking). Furthermore, via *in situ* HCR and time-resolved scRNAseq, we dissect how starting conditions influence the overall cell type composition and the spatial arrangement of germ layers as well as key axial patterning markers: Gastruloids from mESCs cultured in base medium develop *Gata6^+^* and *Eomes^+^* anterior endodermal and cardiac mesodermal regions, while base+1i- and base+2i-derived aggregates lack these lineages, favoring *Sox1^+^* and *Sox2^+^*neuro-ectodermal anterior domains, despite generating axially elongated structures in all cases. The similarity of Hox gene expression coupled with the presence of similar somitic and trunk neuro-ectodermal cell types (although with different proportionalities) in gastruloids across conditions could be the reason why such differences in spatiotemporal patterning have gone unnoticed before. Conversely, timed, but non-localized application of external signals can coax gastruloid from one condition to generate cell types present in another condition. Specifically, we show that upregulation of Activin/Nodal signaling induces anterior (*Eomes^+^*) mesodermal poles in aggregates from base+1i and base+2i, making them resemble base medium-derived gastruloids. This demonstrates the ability to reprogram axial patterning dynamics through external cues. Finally, mixing ESCs from different media conditions enables the generation of gastruloids containing anterior cardiac mesoderm, neuro-ectoderm, and endoderm, showcasing their combinatorial potential for tissue engineering.

Other recent studies further highlight how distinct media compositions, both before and after aggregation, can influence gastruloid developmental dynamics, albeit it should be mentioned that specific effects may vary between cell lines. For instance, mESCs cultured alternatingly in classical (i.e. serum-free) 2i and ES-LIF reproducibly give rise to elongated gastruloids with comparatively higher mesodermal diversity (***40***). Moreover, a comparison of aggregate development in home-made versus commercial N2B27 demonstrates how 3D differentiation medium formulation affects gastruloid morphogenesis and cell fate specification (***55***), with home-made N2B27 favoring larger anterior domains and increased spinal cord-related gene expression.

While the prevailing model of gastruloid development as a surrogate for embryonic scenarios assumes a predefined sequence of differentiation and germ layer emergence, we present evidence that cell type emergence can vary remarkably across conditions yet still lead to robust axial elongation and patterning. Our work hence suggests a broader developmental flexibility underlying these processes. Notably, this extends beyond the evident temporal differences in lineage specification events, whereby naive mESCs take longer to differentiate than primed ones, thus altering T polarization and aggregate elongation dynamics. Such flexibility enables gastruloids to mimic various subsets of embryonic developmental trajectories through external signals or changes in the local environment, e.g. by mixing pluripotent cell populations **(Fig. 5E)**. Similar to how naive pluripotency can be maintained by modulating parallel signaling pathways (***38***), our findings demonstrate that, during differentiation, multiple patterning trajectories resulting in the generation of an elongated body axis are accessible to gastruloids. A key paradigm here is the absence of the native (extra-)embryonic context and associated mechanical as well as geometrical constraints in the gastruloid that allows for such variability in spatial allocation of tissue types along the AP axis *in vitro*. The breadth of the underlying trajectories or, to employ a summarizing term, “alternative developmental modes” available to mammalian PSCs has only recently begun to be studied (***7*,*22***). This will be accompanied by an increasing recognition of the numerous ways and constituent tissue fates through which acquisition of body plan polarity can be achieved in early-branching metazoans (***56***) that should altogether provide new insights into how animal embryonic form can emerge, diversify and evolve.

## Acknowledgements

We thank the Universitat Pompeu Fabra (UPF) genomics core facility and the European Molecular Biology Laboratory (EMBL) Heidelberg GeneCore facility for performing the the sequencing and the Centre for Genomic Regulation (CRG) Protein Technologies Unit for their help with designing and cloning the mESC double reporter allele as well as the CRISPR-Cas9 donor plasmids. We also thank the CRG Tissue Engineering Unit for the preparation of mESC base medium and, in particular, Marta Vila Cejudo, Jacqueline Severino and Laura Batlle Morera for generating and validating the transgenic mESC 2KI reporter line used in this study. We further thank the members of the Trivedi and Ebisuya labs at EMBL Barcelona as well as Alfonso Martinez Arias for critical feedback on the manuscript. We thank the Mesoscopic Imaging Facility (MIF) of EMBL Barcelona for assistance with imaging and microscope procurement.

## Resource availability

### Lead contact

Requests for further information and resources should be directed to and will be fulfilled by the lead contact, Vikas Trivedi.

### Materials availability

The 2KI reporter mESC line (T-p2a-H2B-eGFP and CAG::H2B-mKO2) generated in this study is available upon request from the lead contact. Plasmids generated in this study are available upon request at the CRG Protein Technologies Unit and the CRG Tissue Engineering Unit.

### Data availability

The single-cell transcriptomic data from this study are publicly available on ArrayExpress:

https://www.ebi.ac.uk/biostudies/arrayexpress/studies/E-MTAB-13643

Python and R scripts employed in this study are publicly available on github:

https://git.embl.de/grp-mif/image-analysis/gastruloids_tracking_2024

https://github.com/LabTrivedi/Data-analysis-of-cell-tracks

https://github.com/kerim-a/gastruloid_scRNA_pluripotent-states

https://github.com/kerim-a/TGMM_TrackAnalysis_byNicolaGritti

All other data types will be shared by the lead contract upon request.

## Author contributions

VT: Supervision, Funding acquisition, Resources. VT & KeA: Conceptualization, Project Administration, Methodology, Writing - original draft, Writing - review & editing. KeA: Investigation, Data curation, Formal analysis, Visualization, Validation. FN: Investigation, Formal analysis, Data curation, Resources, Writing - review & editing. NG: Software, Formal analysis, Data curation, Visualization, Writing – review & editing. JFR: Software, Formal analysis, Visualization, Data curation. KrA: Resources, Validation, Writing – review & editing. MPH: Formal analysis, visualization. DO: Supervision, Formal analysis, Visualization, Writing - review & editing. JS: Supervision, Funding acquisition, Resources, Writing - review & editing.

## List of Supplementary Materials

Figures S1 to S6 and legends for Movies S1 to S3.

Movies S1 to S3, all related to Figure S3.

## Materials and Methods

### Mouse ESC culture

Mouse ESCs (mESCs) were cultured as previously described (***57***). In brief, T-p2a-H2B-eGFP and CAG::H2B-mKO2 (2KI) double transgenic reporter mESCs as well as G4 wild-type (WT) mESCs (***58***) were maintained in either base, base+1i or base+2i medium. Base medium consists of KnockOut DMEM (ThermoFisher 10829018), 1x MEM Non-Essential Amino Acids Solution (100x, ThermoFisher 11140050), 1x GlutaMAX (100X, ThermoFisher 35050061), 1x Sodium Pyruvate (100x, ThermoFisher 11360070), 1x Pen/Strep (100x, ThermoFisher 15140122), 50μM B-mercaptoethanol (50mM, ThermoFisher 31350010), 10% fetal bovine serum (FBS) (batch tested) and home-made Leukemia Inhibitory Factor (LIF, also batch tested). For base+1i, 0.3μM (final concentration) of the MEK/ERK inhibitor PD03 (Sigma PZ0162) was added to base medium, deviating from the commonly employed 1μM in order to protect (epi-)genetic integrity of the cultured cells (***43***). For base+2i, we added 1.5μM of the Src-kinase inhibitor CGP77 (Sigma SML0314) and 3μM of the GSK-3B inhibitor CHIR99 (Sigma SML1046) to base medium (both final concentrations).

Base medium preparation and LIF as well as FBS batch testing were conducted by the Tissue Engineering Unit of the Centre for Genomic Regulation (CRG) which can provide plasmids and detailed protocols upon request. LIF-conditioned medium was generated using HEK293T cells transfected with a pCMV_LIF plasmid. Batch validation was achieved by performing the following tests with complete base medium containing the LIF using a E14tg2a mESC line: i) Clonal assay and Alkaline Phosphatase (AP) staining, ii) steady state culture test for 5 passages and assessment for pluripotency by morphology and AP staining. FBS batch testing was performed over 4-6 different commercial batches. Validation was achieved by performing the following tests with complete base medium containing the respective 10% FBS to be assessed using a E14tg2a mESC line: i) Clonal assay and Alkaline Phosphatase (AP) staining, ii) steady state culture test for 5 passages and assessment for pluripotency by morphology and AP staining.

In general, mESCs were cultured on 0.1% gelatin-coated (Millipore, ES-006-B) tissue culture-treated 25cm^2^ flasks (T25 flasks, Corning, 353108) in a humidified incubator (37^∘^C, 5% CO2, Thermo Fisher Scientific). mESCs were passaged every 2-3 days and the culture medium was exchanged for fresh, prewarmed medium on alternating days. Cells were cultured up to 50-70% confluency prior to passaging. After thawing of frozen stocks, mESCs were kept in the respective medium condition (base, base+1i or base+2i) for at least 3 passages before experimental use and the initially chosen medium condition was never changed for a given batch.

### Generation and validation of the double transgenic T reporter mESC line

The 2KI double transgenic reporter mESC line was generated as an integrated CRISPR-Cas9 gene editing project between the Protein Technologies and the Tissue Engineering Units of the CRG (Centre for Genomic Regulation). Hence, more detailed versions of the protocols, including validation data and all plasmids used are available at these facilities upon request. The CRISPR-Cas9 gene editing machinery was designed and assembled by the CRG Protein Technologies Unit.

Transgenesis was performed in a germline competent G4 mESC line (***58***) as a two-step process: First, the CAG::H2B-mKO2 allele was introduced and a stable line was generated, then the T-p2a-H2B-eGFP tag was added into the single transgenic mESCs. To achieve genetic modifications, mESCs were cultured in base medium as described above, however in presence of immortalized mouse fibroblasts feeders (iMEFs, made by the CRG Tissue Engineering Unit), seeded in advance. For genomic DNA extraction for PCR screenings, cells were grown without iMEFs.

For each individual knock-in, 3 sgRNAs were cloned into a PX458 plasmid and sgRNA efficiency was determined via a T7 endonuclease I assay. The best performing sgRNA (Table 1) was then employed for generation of the modification of interest. Thus, for each sgRNA 1×10^6^ G4 mESCs were nucleofected with 10ug of the respective PX458 sgRNA plasmids using a NEPA21 electroporator (Nepagene) and cultured in base medium for 48 hours (h). Thereupon, cultures were stopped, genomic DNA (gDNA) was extracted and amplified by PCR using specific T7 primers. PCRs were conducted as follows: KAPA HiFi HotStart ReadyMix was mixed with primers (0.3μM each), 100ng template DNA and PCR grade water. The reactions were performed with the following settings over 35 cycles: Initial denaturation: 95^∘^C, 3min; Denaturation: 98^∘^C, 20sec; Primer annealing: 63.2^∘^C, 15sec; Extension: 72^∘^C, 2min; Final extension: 72^∘^C, 5min. PCR products were purified using a MinElute PCR Purification Kit (QIAGEN) and employed in T7 endonuclease reactions by mixing 100ng PCR template with buffer (NEB, New England Biolabs) and 0.5μl T7 endonuclease. Results were visualized via a 2% agarose gel.

**Table 1.**
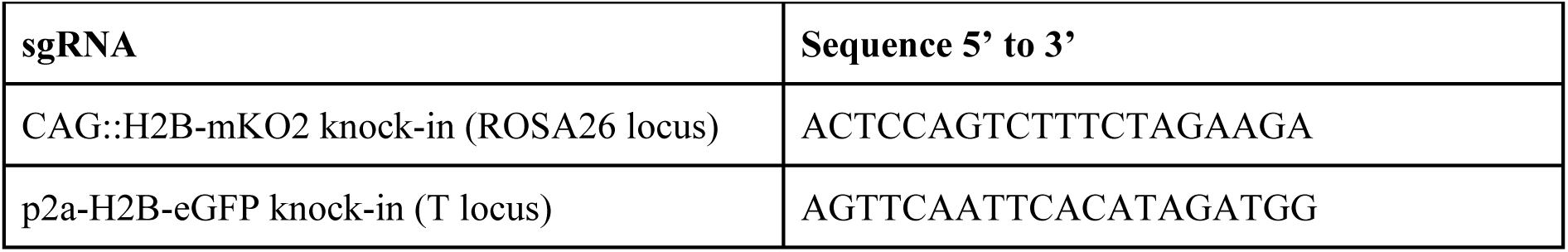
List of sgRNAs.

CRISPR-Cas9 mediated knock-in generation was achieved by nucleofecting 1×10^6^ G4 mESCs with 5μg PX458 sgRNA plasmid and 5ug homologous recombination and knock-in donor plasmid using a NEPA21 electroporator. Nucleofected cells were then plated and cultured in base medium + iMEFs and further treated with Neomycin at 400μg/ml during 3 days. After recovery from nucleofection, for generation of clones, single cells were sorted into flat-bottom 96WP (Thermo Scientific, 167008) via a BD Influx Flow Cytometer with the following parameters: Nozzle: 100μm; Pressure: 12psi; Sample differential: 12 psi; Sorting by FSC/W and SSC/W. Cells were then grown in base medium + iMEFs. Clones were then expanded into a working plate and a replicate plate. From the replicate plate, gDNA was extracted and PCR-based screening (see above for settings) was performed to detect clones containing the knock-in of interest. From the working plate, suitable clones were expanded in flat-bottom 6WPs (Corning, 353046), cryopreserved and further evaluated by PCR (Table 2). Finally up to 4-5 validated clones were analyzed by Sanger sequencing to validate the expected insertion of the modification of interest. In case of the T-p2a-H2B-eGFP reporter knock-in, reporter expression was confirmed via a mesodermal differentiation assay, including 48-72h culture in N2B27 with 3μM CHIR99 and 50ng/ml Activin A (R&D Systems, 338-AC-010), and subsequent imaging on a Opera Phenix HCS system (PerkinElmer).

**Table 2.**
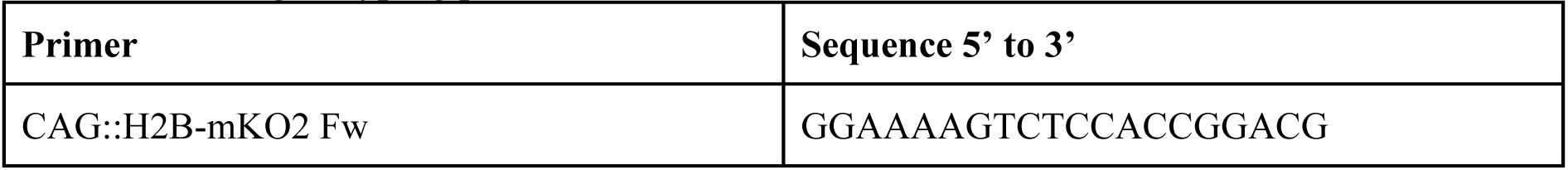
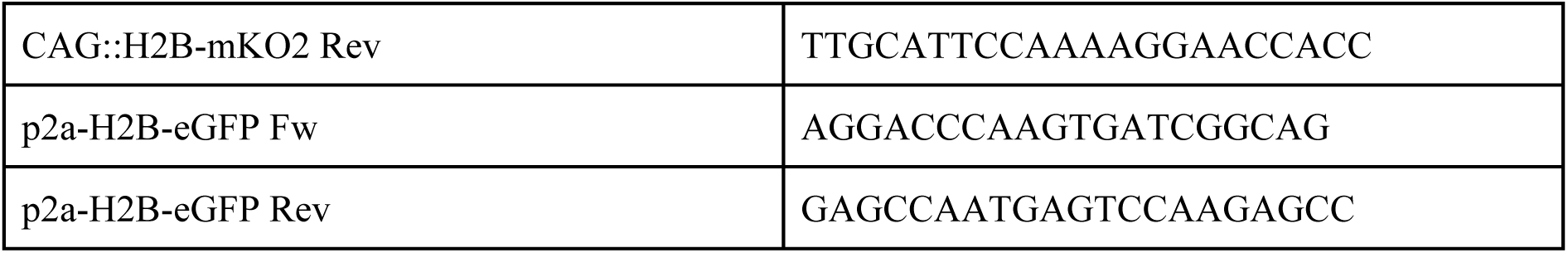
List of genotyping primers:

### Gastruloid generation

Gastruloids were generated as previously described (***57***). To sum up, mESCs growing in base, base+1i or base+2i medium were gently rinsed with 5ml pre-warmed DPBS+/+ (DPBS with Mg and CaCl2, Sigma, D8662). DPBS^+/+^ was replaced with 1ml pre-warmed TrypLE Express (Gibco, 12604013) and the cells were incubated at 37^∘^C for 1-2min. Thereupon, TrypLE was diluted via application of 4ml base medium. Cells were collected via gentle pipetting, the suspension was transferred to a 15ml centrifuge tube and centrifuged for 3min at about 180g (900-1000rpm). Then the supernatant was aspirated and cells were resuspended in 10ml warm DPBS^+/+^ (washing). This step was repeated and after another centrifugation round (3min, 180g), the pellet was resuspended in 0.5ml to 1.5ml of pre-warmed differentiation medium N2B27 (Ndiff 227, Takara, Y40002) via pipetting up-and-down 5-15x with a P1000 pipette until remaining clumps of mESCs were separated into single cells.

Hereafter, mESCs were counted using an automated cell counter (Countess II, Invitrogen). For this, 10μl of cell suspension were mixed with 10μl staining mix (Invitrogen, T10282) and loaded on a counting slide (Invitrogen, C10228). The calculated volume of the cell suspension was then added to the required amount of warm N2B27 and transferred to a sterile reservoir. Using a microchannel pipette, 40μl of plating suspension were added to each well of a U-bottom, low adhesion 96-well-plate (96WP, Greiner, 650970) which was thereafter returned to the incubator and maintained at 37^∘^C and 5% CO2. For gastruloids from mESCs grown in base medium, we seeded 300-350 cells per well of a 96WP (cells/well), for base+1i, we seeded 400 cells/well, for base+2i, we seeded 500 cells/well. After 48h, further 150μl of prewarmed N2B27 containing 3μM CHIR99 (Sigma, SML1046) were pipetted into each well, followed by daily medium exchange with just N2B27 if gastruloids were grown beyond 72h. In case source mESCs were grown in base+2i medium, the CHIR99 pulse was applied between 72-96h post-aggregation. For anterior mesoderm upregulation experiments (see Fig. 5), 50ng/ml Activin A (R&D Systems 338-AC-010), was co-applied with CHIR99.

### 2D Imaging and quantitative analysis of gastruloids

Cell aggregates cultured in round-bottom 96WPs were imaged with an Opera Phenix High Content Screening (HCS) System system (PerkinElmer) in wide-field mode using a 10X air objective (NA 0.3). Focus heights were manually adjusted for a given plate, depending on size of the gastruloids. For time lapse experiments, the incubator module was set at 37^∘^C and 5% CO2. Medium evaporation was prevented by replacing the lid of the round-bottom 96WP with a MicroClime environmental microplate lid (Labcyte, LLS-0310) filled up with DPBS-/- to maintain humidity. Images were then acquired with a 10min interval.

After acquisition, images were compiled in single multichannel files. All images were segmented, analyzed and plots were generated via MOrgAna, a machine learning (ML) pipeline implemented with a custom-written, open source Python code (***59***). Segmentation masks (and midlines) for each gastruloid at every timepoint were thus generated as previously described (***7*,*59***). Quantifications of images acquired on the Opera Phenix system generally represent data from one experimental/technical replicate, encompassing numerous biological replicates, several conditions, usually across consecutive developmental days. Results were corroborated by performing (at least) one more experimental replicate.

#### Eccentricity

To account for the bending of gastruloids during elongation, eccentricity was computed on a computationally straightened version of the final mask. Briefly, we used distance transform to find the midline and the width of the gastruloid, and computed eccentricity according to 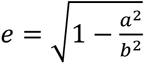, here a and b represent the major and minor axis length of the ellipse with the same second moments as the straightened gastruloid mask.

#### AP fluorescence intensity profiles

To compute the AP fluorescence intensity profiles, for every position along the midline of the gastruloid, the average pixel intensity in the fluorescence image channel along the direction orthogonal to the midline was computed. The AP profile was then oriented such that the posterior part of the gastruloid always represented the highest T expressing pole. For analysis across conditions, the gastruloid length was normalized to values between 0 (anterior) and 1 (posterior). T fluorescence values were normalized globally (i.e. using all datasets included in the respective set of plots). Since the employed N2B27 medium exhibits auto-fluorescence upon exposure to the 488nm laser, for each plot, background subtraction was performed by subtracting the intensity values outside the gastruloid mask (“Background subtraction type: Background” in the MOrgAna interface).

#### Definition of a polarization index

We defined a system to represent polarization of a gastruloid based on the fluorescence expression over the fitting midline over its main axis. To do so, we computed the integral of the fluorescence over the relative midline length (0 at one point and 1 at the other) and we define p_*_ as the point that splits the integral in half. We then took the maximum between p∗ and 1− p_*_, so 0.5 ≤ p_*_ < 1. When p_*_ = 0.5, this means that the point that splits the integral in half is the midpoint, which means that we have a symmetrical distribution and there is no polarization. On the other hand, the more polarized the gastruloid is, the larger will be p_*_. However, we remark that 1 is a theoretical limit, so the maximum value that p_*_ can have is p_max_ < 1. From previous computations, we can define two measures for the polarization. We call the first one the absolute polarization p_abs_ which linearly maps p_*_ from [0.5, 1] to [0, 1]. This measure will never reach 1, but it is appropriate to compare polarization across different conditions. The second measure is the relative polarization p_rel_ and maps p_*_ from [0.5, p_max_] to [0, 1]. By construction, p_rel_ can be equal to one, which is more appropriate to study the changes of dynamics within tracks belonging to the same condition.

#### Computation of normalizing parameters for hexbin plots

For the hexbin plots, there are three variables that are normalized: area, midline length and average intensity. Also, for each variable, there is a different normalizing parameter, depending on the condition. The normalization is done as follows:

- Area and midline length: We take the data over the earliest time, for the gastruloids that express T from a particular condition. Then, we compute the average of the early gastruloids area or midline length.
- Average intensity: We take the data over the earliest time, for the gastruloids that do not express T from a particular condition. Then, we compute the average of the early gastruloids’ averaged intensity.

### Aggregate fusion assays and analysis

For imaging homotypic aggregate fusion, at 24hpa two aggregates from mESCs of a given medium condition (base, base+1i or base+2i) were placed in a single well of a 96WP using cut-off (UV-sterilized scissors) P200 tips. In order to generate 24hpa gastruloids at roughly equal size which is crucial for comparability of fusion dynamics and inferred parameters, we aggregated 500 cells/well for base medium, 350 cells/well for base+1i and 300 cells/well for base+2i. Live imaging was performed with an Opera Phenix High Content Screening (HCS) System system (PerkinElmer) as described above. Images were acquired with a 10min interval. Segmentation masks of the fusing aggregates were generated via MOrgAna (***59***). These were used for further analysis of fusion dynamics and parameter inference, according to (***44,45***), however differential aggregate growth was also accounted for. To achieve this, the evolution of aggregate radius was quantified for the time period of the fusion assay for 2 individually developing aggregates from each medium condition. A detailed explanation of the employed continuum model for the fusion of two proliferation aggregates and the inference of material properties can be found on github: https://github.com/kerim-a/aggregate-fusion-analysis/tree/main

### In situ Hybridization Chain Reaction (HCR)

Gastruloids were harvested from U-bottom 96WPs in UV-sterilized organoid collection plates (OCPs, patent EP3404092A1) and transferred to 2ml Eppendorf tubes using RNase free wide-bore P1000 pipette tips. After 3 washes for 5min each with 1ml DPBS^-/-^ (DPBS without Mg and CaCl2, Sigma D8537), samples were fixed overnight (ON) at 4^∘^C in 1ml 4% PFA in DPBS^-/-^. The following day, samples were dehydrated by 3 DPBS^-/-^ washes (5min each, room temperature), 4 washes in 100% Methanol (MeOH) for 10min each at room temperature (RT) and 1 wash in 100% MeOH for 50min at RT. Hereafter, samples were stored at −20C, if required.

Rehydration was performed through a series of sequential washes at RT: 5min in 75% MeOH in DPBS^-/-^ with 0.1% Tween (PBST2), 5min in 50% MeOH in PBST2, 5min in 25% MeOH in PBST2 and 5x in PBST2 for 5min each. Samples were then transferred to a 1.5ml Eppendorf tube and 500μl probe hybridization buffer (PHB, Molecular Instruments, HCR v3.0) was applied for 30min at 37^∘^C. Immediately afterwards, probe solution was prepared by adding 2pmol of each probe of interest (all probes were ordered through Molecular Instruments for HCR v3.0) into 500μl PHB and maintained at 37^∘^C for 30min. Thereupon, probe solution was applied to the samples, replacing the PHB, and incubation was performed ON at 37^∘^C.

The next day, 4 washes (15min each, 37^∘^C) were conducted in 500μl probe wash buffer (PWB, Molecular Instruments, HCR v3.0) preheated to 37^∘^C and 2 washes (5min each, RT) were performed in 5x SSCT (5x sodium chloride sodium citrate, 0.1% Tween in ultrapure H2O). Hereafter, amplification buffer (Molecular Instruments, HCR v3.0) was equilibrated at RT and SSCT was replaced with 500μl amplification buffer (equilibrated at RT), followed by incubation for 30min at RT. 30pmol of each hairpin (h1 and h2, Molecular Instruments, HCR v3.0) were then separately aliquoted (2μl of 3μM stock), heated to 95C for 90s and cooled for 30min in the dark at RT. Thereupon, hairpins were mixed in 500μl amplification buffer (AB), pre-equilibrated at RT, to a final concentration of 60nM per hairpin. AB was removed from samples and replaced with hairpins in AB, followed by incubation ON at RT, in the dark.

Hereafter, 5 washes in SSCT were performed at RT: 2x 5min, 2x 30min and 1x 5min. Samples were then optionally stored at 4^∘^C in the dark for 24h to 2 weeks. Gastruloids were imaged in a flat-bottom CellCarrier-96 ultra plate (PerkinElmer, 6007008) on an Opera Phenix HCS system (PerkinElmer) in the confocal mode, using a 20x water objective. We note that single, representative gastruloids were cropped from the original images and rotated using FIJI/ImageJ (***60***) for display purposes. This is because samples were imaged in random orientations due to the Opera Phenix HCS not allowing manual stage movement, leading to many samples being distributed along the edge of the field-of-view.

#### HCR fluorescence intensity profiles

For multi-channel AP fluorescence intensity analysis of *in situ* HCR data, a custom add-on python script for MOrgAna was employed after generating sum intensity projections of HCR imaging data in FIJI and calculating masks as well as AP fluorescence intensity separately for each channel as previously described. Briefly, the T channel was used as the reference to delimit the AP axis and fluorescence intensity from the other channels was plotted along this thereby determined axis. Fluorescence intensity for all channels were normalized to their own respective maxima across all included developmental timepoints.

### Light sheet fluorescence microscopy (LSFM) imaging of gastruloids

For LSFM imaging, mosaic gastruloids were generated from 5-10% of 2KI G4 mESCs and 90-95% of G4 WT mESCs were grown in 96WPs as described above. At the start of the designated imaging window, aggregates were transferred into a Viventis LS1 live system sample holding chamber filled with 0.5ml N2B27 containing 1x Pen/Strep (Gibco, 15140122) (N2B27PS) using a P200 pipette tip cut off with UV-sterilized scissors. 1ml N2B27PS was then carefully added and the chamber was mounted on the LS1 live system for imaging with the temperature and CO2 modules set to 38-39^∘^C (achieving 36.5-37^∘^C water temperature in the sample holding chamber) and 5% CO2, respectively. Imaging was performed with a Nikon 25XW NA 1.1 water immersion objective and images were captured on an Andor Zyla 4.2 sCMOS camera with a time-interval of 5min across ∼8.5-16.5h long time windows.

### LSFM time-lapse data registration and cell tracking

The images containing the nuclear signal of the 4D datasets were analyzed to extract single cell trajectories using TGMM (***61***). Because gastruloids are freely floating in the culture media, their positions in two consecutive images might change, and their cumulative changes over time make the task of tracking single cells over time particularly challenging. For this reason, before processing the 4D datasets with TGMM, we performed computational registration of the images:

First, we generated an isotropically resolved stack and detected most of the cell positions by applying a Gaussian filter with a sigma equal to the approximate radius of single cells and finding all the local maxima within the 3D stack. To avoid detecting maxima in dark regions of the images, maxima corresponding to pixel values lower than 200 were ignored.

Next, for each pair of consecutive timepoints T and T+1, we determined a 4×4 rigid transformation M consisting of a rotation R and a translation T:

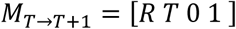

and applied it to the cloud points corresponding to the cell positions at time point T:

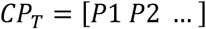

The transformed cloud point *CP′*_*T*_ was then overlaid to the target cloud point at T+1, and the distances between neighboring cells were computed. The optimal transformation 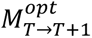 was obtained using a gradient-based minimization algorithm available in the scipy Python library. We therefore obtained a sequence of transformations 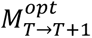 that optimally registered consecutive cloud points *CP_T_* and *CP_T_*_+1_. The global transformation to register cloud points from any arbitrary time point was then obtained by standard matrix multiplication of the 4×4 instantaneous transformations:

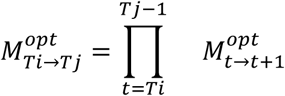

#### Such that

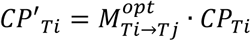

Was optimally registered with the cloud point *CP_Tj_*.

Finally, the 3D image stacks were computationally registered by applying the resulting rigid transformation to the isotropically resolved stack. To avoid computer memory issues while running TGMM and to speed up computation, registered image stacks were rendered anisotropic with the same voxel size as the input images.

The registered images were subsequently processed using a top hat transform, defined as the difference between the input image and its morphological opening by a structuring element. To enhance cell detection, we chose a disk with size of approximately two cell diameters as structuring element. The resulting images displayed reduced scattering effects and more homogeneous signal-to-noise ratio (SNR), thus facilitating the cell detection task of TGMM.

Registered datasets were then processed with TGMM, comprising nuclei segmentation and tracking. Main parameters (background threshold and segmentation τ-level) were adjusted for each sample and advanced parameters were left as default. After converting the image dataset into xml/HDF5 format for visualization in the FIJI plugin BigDataViewer, TGMM output tracks were imported into Mastodon (linked to the xml/HDF5 file), a FIJI plugin designed for manual curation and feature extraction of cell tracks. We thereupon imported the resulting computed cell and track features (via Mastodon, all available parameters) in Python, using custom written Jupyter Notebooks, and removed untracked cells, that is, cells that appear in one time point but do not belong to any track.

#### LSFM data analysis

##### Criteria for fluorescence threshold

Given the tracks of cells over time, we define a criteria with which to compute two thresholds: T^-^ and T^+^. Values between T^-^ and T+ were treated as a transition interval. To define the thresholds, we plotted both the normal and accumulated histogram of all fluorescence values over all the tracks we have in a period of time. The threshold T^-^ is taken as the inflection point of the accumulated histogram. After that, we assume that the normal histogram follows a Poisson distribution Pois(λ), where λ is the mean of all fluorescence, and we compute its variance. The upper threshold T^+^ is defined then as T^+^ = T^-^ + 5Var[Pois(λ)].

##### Mean Squared Displacement (MSD) analysis

To study the behavior of cells provided their tracks, we consider the MSD computation of a track x_i_(t), given by:

MSDi(τ) = < |x_i_(t + τ) - x_i_(t)|^2^ >, where < · > represents the average over all frames t for x_i_(t).

In order to know the type of cell dynamic, we study the log-log of MSD(τ) in relation to τ. If the best-fitting slope D of the log-log is equal to 1, the system is diffusive. If it’s equal to 2, it is ballistic. Values D < 1 represent confined space.

##### Categories for T expression transition dynamics

To establish the criteria of T expression evolution, we take the average between the threshold for T^-^ and T^+^, so there is only one threshold that splits between the T^-^ and T^+^ side. The five categories that result to classify T transition dynamics are:

- Pure T^-^: Cell expression remains at the negative side along over all its tracking, up to two time frames.
- Pure T^+^: Cell expression remains at the positive side along over all its tracking, up to two time frames.
- T^-^ to T^+^: Cell expression begins at the negative side and switches at some point to the positive one. Then, it remains there for up to two time frames.
- T^+^ to T^-^: Cell expression begins at the positive side and switches at some point to the negative one. Then, it remains there for up to two time frames.
- Fluctuating: Any cell that does not fit the previous categories.

##### Track visualization

We set a threshold where we consider only cells tracked over at least 5 hours. Then, given these cells, we plot the vectors from initial to final position, centering them at the origin of axis (0,0). Plots are split over the categories described above. In each plot, all vectors are plotted, but only the ones matching the category are colored, while the rest are in transparent grayscale.

### Preparation of gastruloids for scRNA-seq

To ensure sufficient numbers of dissociated cells for each experimental condition, 3-4 U-bottom 96WPs of gastruloids were generated for the pre-CHIR99 pulse timepoint, 2-3 for the post-CHIR99 pulse timepoint and 1 for the maximum elongation timepoint. For the 0hpa timepoint, mESCs were grown in a T25 flask as described above. 96WPs were either harvested using UV-sterilized OCPs or via manual pipetting directly from the 96WP with an RNAse-free wide-bore P1000 tip (Thermo Fisher Scientific, 2079G). Individual aggregates were thus pooled into a 15ml centrifuge tube (Corning 430791). When all plates from a given condition were harvested, the centrifuge tube was placed into an incubator (37^∘^C, 5% CO_2_) and harvesting proceeded with plates from the next condition.

Once all samples were collected, thus yielding one 15ml centrifuge tube per condition, aggregates were centrifuged at 180g for 2min. Hereafter, the supernatant was removed and 1ml TrypLE (Thermo Fisher Scientific, 12604) was added per tube for dissociation and samples were incubated for 5min (at 37^∘^C, 5% CO2). Then, gentle pipetting was performed (2-3x with the P1000 pipette) in order to dissociate aggregates into single cells. TrypLE was diluted using 4ml cold (4^∘^C) 0.1% RNase-free BSA (Thermo Fisher Scientific, AM2616) in DPBS^-/-^ (BPBS) and gastruloids were centrifuged for 2min at 180g. From this step onwards, samples were maintained at 4^∘^C. Hereafter, two washes were performed, each one via removal of the supernatant, addition of 5ml cold BPBS and centrifugation (2min, 180g). Following removal of the supernatant, the pellet was resuspended in 50-500μl cold BPBS (depending on pellet size) by gently pipetting up and down 2-3x with a P200-1000 tip (depending on resuspension volume). The cell suspension was then filtered through a 35μm filtered cap tube (Corning, 352235). Cells were counted as previously described, aiming for a concentration of around 1×10^6^ cells/ml. In case of high concentrations (>2×10^6^ cells/ml), the cell suspension was diluted with cold BPBS and re-counted. For the 0h timepoint, mESCs were detached from the T25 culture flask using 1ml TrypLE as described in the maintenance and splitting protocol. Following dilution (4ml base medium) and centrifugation (180g, 3min), the supernatant was discarded and cells were washed with 5ml cold BPBS. Cells were then kept on ice, until samples from other experimental conditions reached this washing step. At this point, all samples were joined together for centrifugation and further procedures illustrated above.

### Library preparation and sequencing

After the generation of single-cell suspensions in BPBS as described above, cells in each sample were barcoded with a Chromium Single Cell 3’ GEM v3.1, a Library & Gel Bead Kit v3.1 (10x Genomics) and a Chromium Single Cell G Chip Kit (10x Genomics) on a Chromium Controller (10x Genomics), followed by cDNA library construction according to manufacturer’s instructions. These libraries were sequenced via a NextSeq 2000 system (Illumina). We read 28, 10, 10 and 90-base pairs for 10x Barcodes with unique molecular identifiers (UMIs), i7 and i5 indices and fragmented cDNA, respectively.

### scRNA-seq data analysis

Quality control (QC), alignment to the mouse genome (mm10), and counting of the sequence reads were conducted with CellRanger v3.1.0 (10x Genomics) to generate feature-barcode matrices. Further QC steps, data normalization, identification of most variable features and dataset integration were performed with Seurat version 4.3.0 (***62***) loaded into R version 4.2.2 (***63***). Low quality cells were removed with the thresholds shown in Table 3.

**Table 3.**
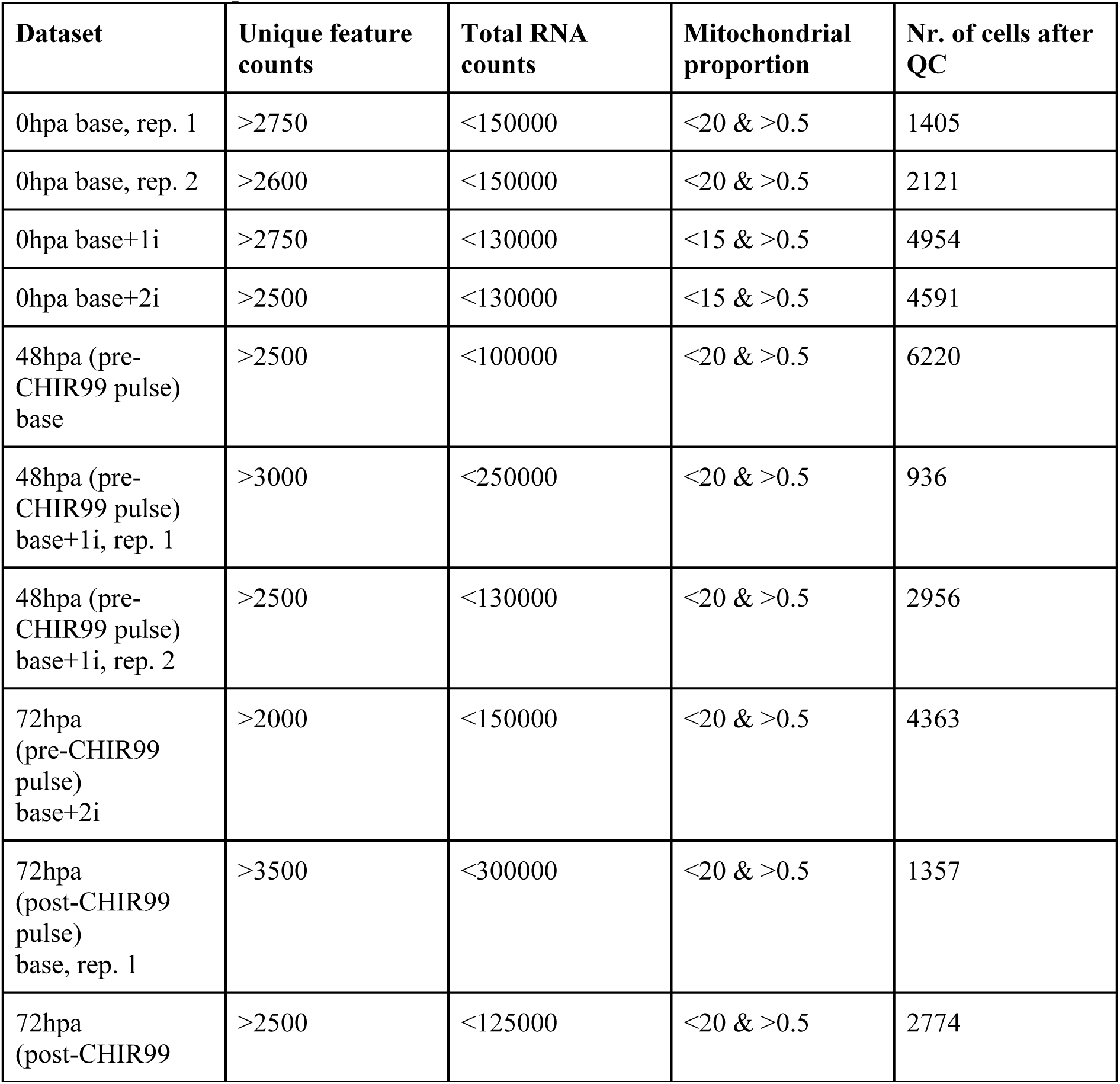
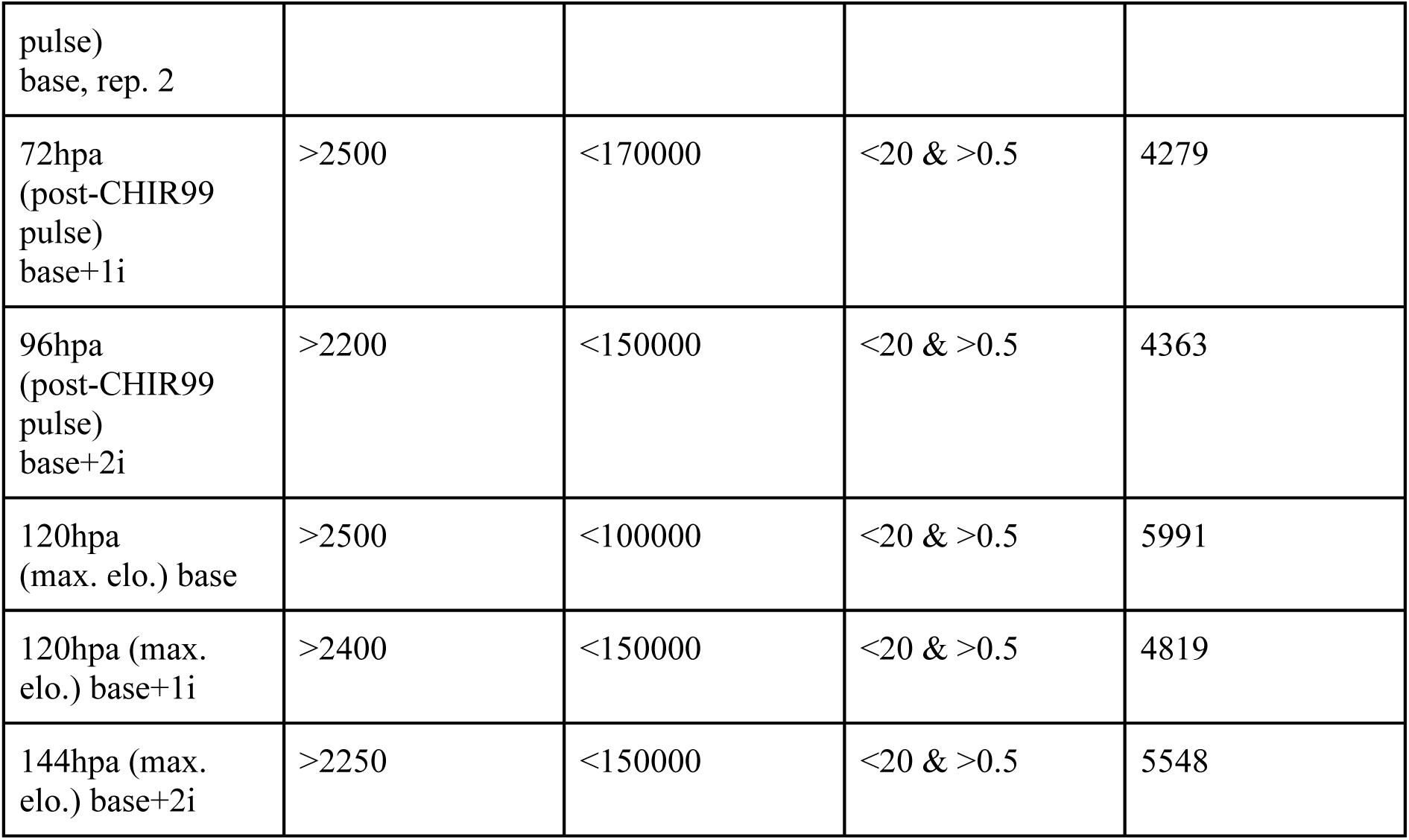
scRNA-seq dataset overview and QC information.

Following this filtering, individual datasets were merged into a single Seurat object (albeit 0hpa/mESC datasets were merged and further processed separately) using the “merge()” function. Normalization via “NormalizeData()” and identification of the most differentially expressed genes via “FindVariableFeatures(, selection.method = “vst”, nfeatures = 3000)” was then performed.

#### Dataset integration, clustering and gene expression analysis

For batch effect correction and dataset integration, we opted for a fast mutual nearest neighbors (fastMNN)-based approach (***64***). To achieve this, 3000 integration features were computed on a list comprising all the individual datasets from the merged seurat object via “SelectIntegrationFeatures(, nfeatures=3000)”. In order to diminish the effect of distinct cell cycle states on the following integration, we removed known cell cycle genes from this list, namely the mouse orthologues from two (integrated) Seurat lists. Thereupon, dataset integration was performed through the “RunFastMNN()” function from SeuratWrappers (version 0.3.1), using the previously computed integration features (minus the cell cycle genes). Clustering was then performed via the “RunUMAP(, reduction = “mnn”)”, “FindNeighbors(, reduction = “mnn”)” and “FindClusters()” functions. In case of the mESC datasets, we set “dims = 1:25” and for the gastruloid data “dims = 1:50” in the former two commands. “FindClusters()” was run with “resolution = 0.3” for the mESC integration and “resolution = 0.7” for the gastruloid integration. Differentially expressed cluster or condition marker genes were identified through the “FindAllMarkers()” and individual clusters were compared using “FindMarkers()”, setting “min.pct = 0.5” and “logfc.threshold” to 0.5 or 0.25 (in case no genes were found with 0.5). Gene expression dotplots were generated via the “DotPlot(, scale.by = “radius”, scale = TRUE)” function. For the hox gene analysis, we set “(…,scale = FALSE)”.

#### Biased cluster identification

In order to identify “biased” clusters, i.e. clusters with elevated proportions of cells from a certain medium condition, for each cluster, we normalized the number of cells from each of the three initial media conditions by multiplication with a scaling factor (total *n* of cells in dataset divided by the total *n* of cells in dataset from a given medium condition). This was performed to account for the conditions not having equal total numbers of cells across the entire integrated dataset(s). Thereby, properly adjusted ratios were computed for each cluster using the adjusted number of cells from a given medium condition in that cluster.

#### Gene expression-based germ layer sorting

For each sequenced timepoint (i.e. 0h, pre-CHIR99, post-CHIR99 and max. elongation), all cells of individual datasets (filtered and normalized as described above) from all three media conditions belonging to the respective timepoint were assessed and sorted according to the following sets of genes.

##### Pre-CHIR99 Mesoderm

Log-normalized expression > 0.2 for any of the following markers: *Ald1ha2*, *Meox1*, *Tbx6*, *Hand1*, *Hand2*, *Kdr*, *Cdx2*, *Eomes*, *T, Meis1*, *Lhx1*, *Bicc1*

##### Post-CHIR99 & Max. Elongation Mesoderm

Log-normalized expression > 0.1 for any of the following markers: *Ald1ha2, Meox1, Tbx6, Hand1, Hand2, Kdr, Cdx2, Eomes,T, Meis1, Lhx1, Bicc1*

##### Pre-CHIR99, Post-CHIR99 & Max. Elongation Ectoderm 1

*Sox1* > 0.2

##### Pre-CHIR99, Post-CHIR99 & Max. Elongation Ectoderm 2

*Epha5* > 0.2 & *Ncam1* > 0.5 & *Sox2* > 0.2

##### Pre-CHIR99, Post-CHIR99 & Max. Elongation Ectoderm 3

*Hes6 > 1 & Tubb3 > 1*

##### Pre-CHIR99, Post-CHIR99 & Max. Elongation Endoderm 1

*Spink1* > 0.2 & *Cldn6* > 0.05

##### Pre-CHIR99, Post-CHIR99 & Max. Elongation Endoderm 2

*Sox17* > 0.1 & *Cldn6* > 0.05

##### Pre-CHIR99, Post-CHIR99 & Max. Elongation Undifferentiated

*Zfp42* > 0.1 OR *Dppa5a* > 0.5

Each cell was hence assigned a germ layer/cell state identity with the above filters applied in the following order: 1) Mesoderm, 2) Ectoderm (any of the 3 filter sets), 3) Endoderm (any of the 2 filter sets), 4) Undifferentiated. Hence, if a cell was designated as “Mesoderm”, but also fulfilled any of the later filters, the “Mesoderm” identity would be overridden and only the latest applicable identity remained. This was done because particularly endodermal and undifferentiated cells can easily be distinguished via few, rather exclusively expressed marker genes, whereas e.g. “Mesoderm” is a comparatively diverse cell state. Thereupon, remaining unsorted or “Undefined” cells from each dataset (<5%) were assessed via gene expression heatmaps (“DoHeatmap()”), highlighting the 5-10 most differentially expressed genes. Based on visual inspection, remaining unlabeled clusters were then optionally allocated to any of the previously defined categories.

## Supplementary Figures

**Fig. S1.**
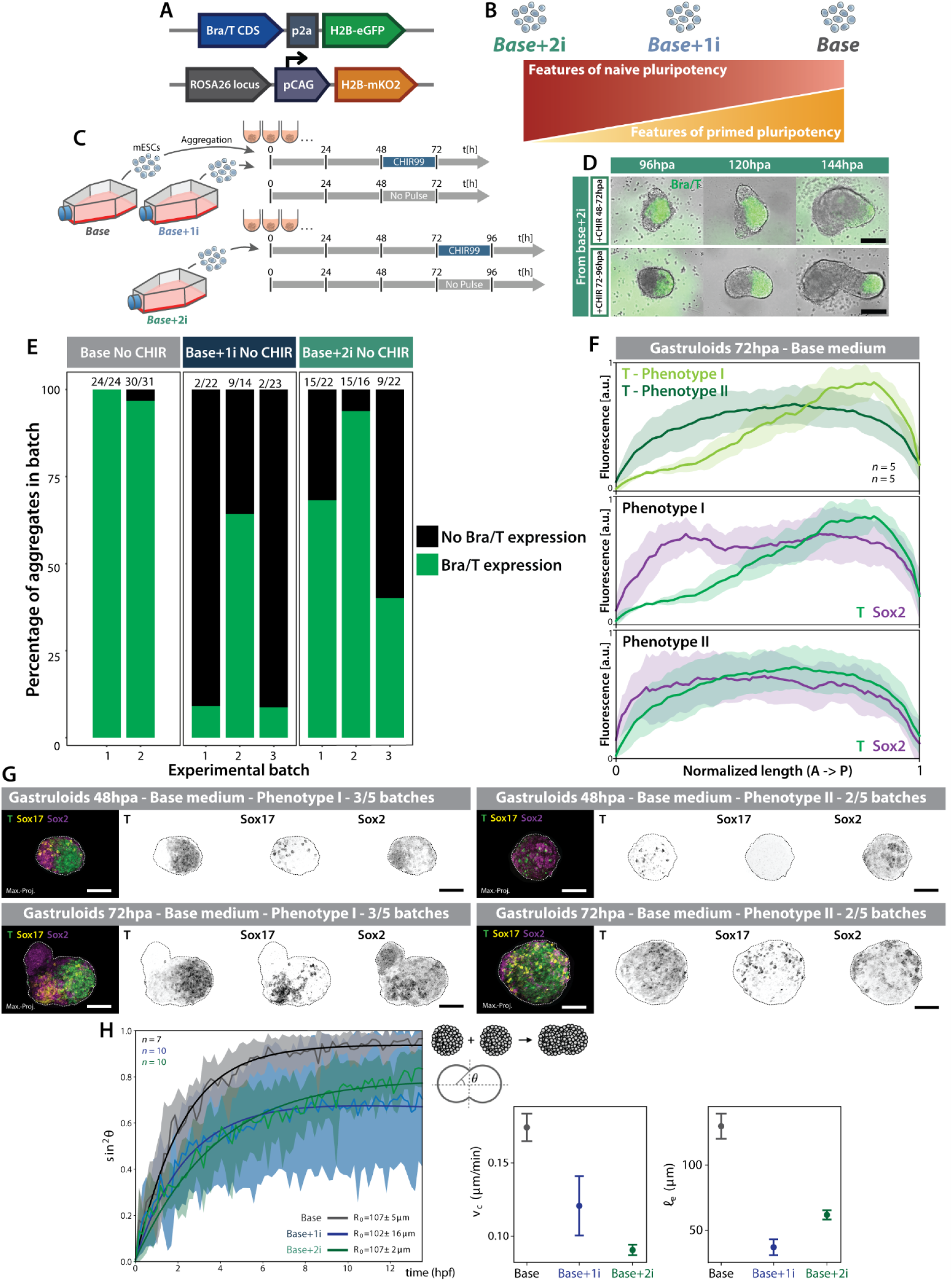
Experimental approach and variability as a result of initial mESC medium condition. **A)** Schematic of the fluorescent reporter alleles from the 2KI double reporter mESC line generated and employed in this study. CDS = coding sequence. pCAG = CAG promoter. **B)** Schematic (adapted from Supp. Ref. 1) describing pluripotent states corresponding to the respective culture media conditions of the mESC populations employed in this study. **C)** Overview of the experimental approach for gastruloid generation from each respective pluripotent state. **D)** Representative images of aggregates from mESCs grown in base+2i, showing that these require a CHIR99 pulse between 72-96hpa to develop a stereotypic, elongated gastruloid morphology. Scale bars = 200μm. **E)** Bar plots showing the percentage of gastruloids in a given batch and from a given source mESC pluripotent state/media condition developing discernible T expression in absence of the CHIR99 pulse. **F)** Quantifications of fluorescence intensity of marker genes from HCR stainings in **G)** along the aggregate’s AP axis. Central lines denote the average across replicated, surrounding shades the standard deviation. Quantifications are based on sum intensity projections of the HCR data. **G)** Representative images of HCR stainings of gastruloids from mESCs cultured in base medium illustrating differences in developmental timing (Phenotype I vs. Phenotype II) between distinct experimental batches. Scale bars = 100μm. **H)** Quantification of 24hpa gastruloid fusion dynamics across different media conditions showing the averaged time evolution of sin^2^ θ, where θ is the fusion angle of the assembly (**43**, also see schematic on top right taken from **44**). Shaded area denotes standard deviation around the mean experimental curve. Solid lines are numerical fits, obtained according to (**43**). R_0_ is the aggregate mean radius. Plots to the right are quantifications of viscocapillary velocity (ν_c_) and shear elastocapillary length (ℓe), inferred from fusion kinetics for each medium condition. See Materials & Methods for details.

**Fig. S2.**
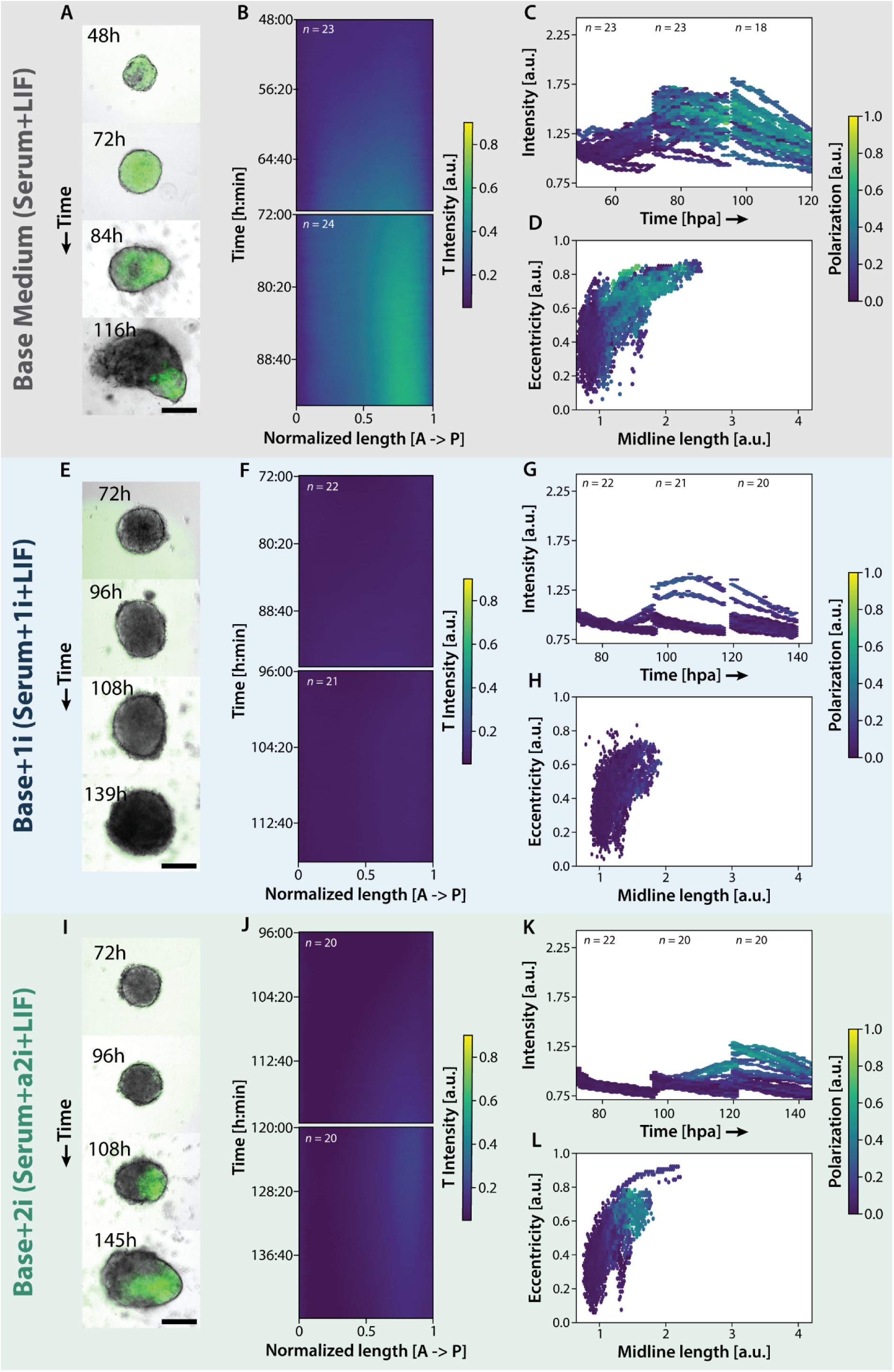
Initial medium condition influences T polarization and aggregate elongation dynamics II. **A), E), I)** Representative images of developing gastruloids grown without a pulse of Wnt agonist CHIR99, from mESCs cultured in either base medium, base+1i or base+2i. Scale bars = 200μm. **B), F), J)** Kymographs displaying average T expression intensity over time along the normalized major or AP axis of the gastruloids (as denoted by T expression) from each medium condition (or pluripotent state). **C), G), K)** Hexbin (2D histogram) plots of individual gastruloid T intensity over developmental time for each condition. The color code indicates the degree of (T) polarization. **D), H), L)** Hexbin plots of individual aggregate eccentricity (describing elongation) over midline length for each condition.

**Fig. S3.**
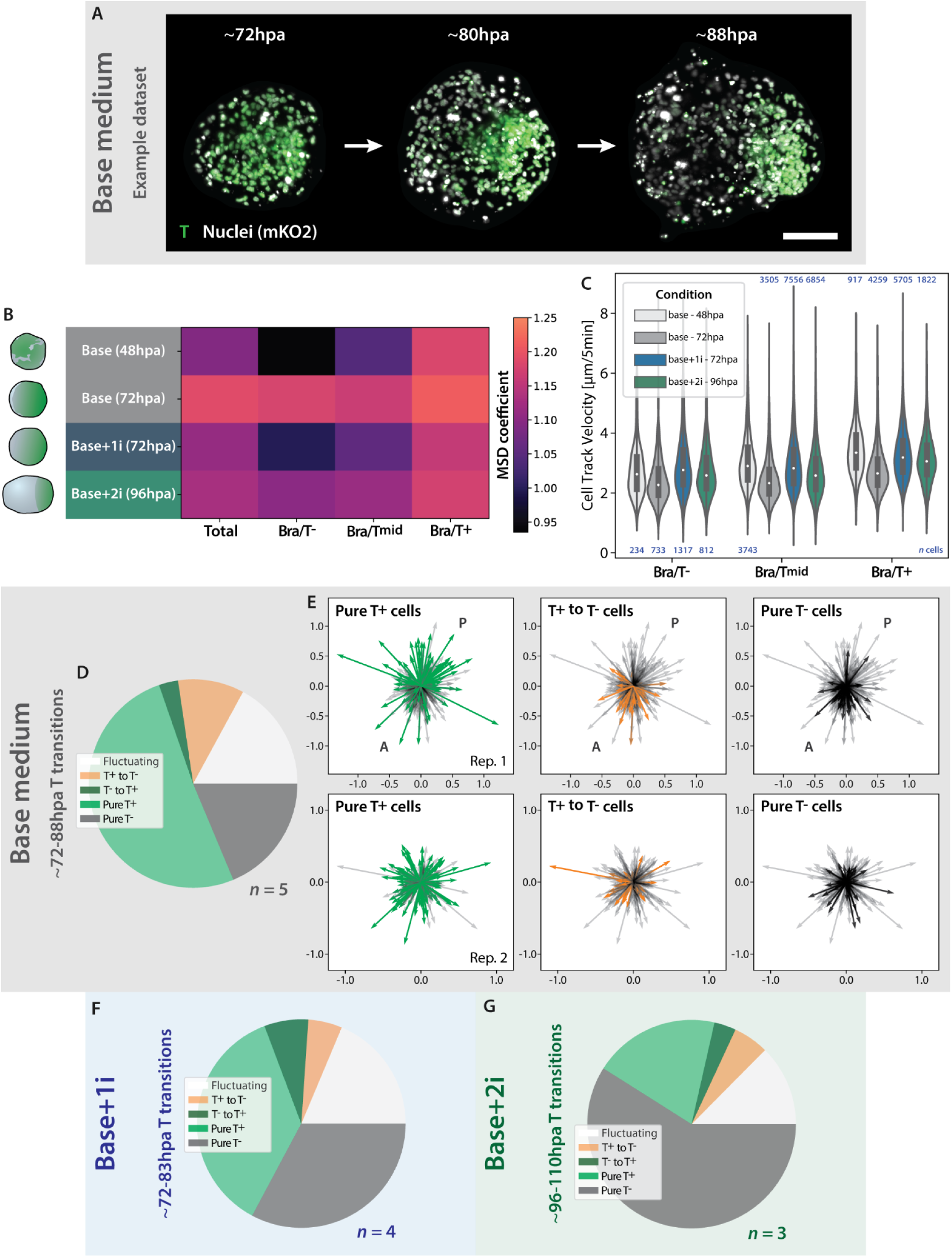
In toto 3D cell tracking of developing aggregates. **A)** Maximum intensity projections of example LSFM dataset (base medium) tracked over ∼16h showing T polarization. Scale bar = 100μm. **B)** Heatmap showing the average MSD coefficient (describing migratory properties) for either all aggregate cells or either the T^-^, T^mid^ or T^+^ populations across source mESC media conditions and imaging time windows. The indicated time describes the onset of imaging which lasted ∼12h in total. Sketches on the left describe gastruloid developmental stage during the respective time window. **C)** Violin plots summarizing the migration speeds of all individual cells, tracked for at least 7 consecutive timepoints, for each respective medium condition and time window. **D)** Pie chart illustrating the proportions of T state transition dynamics and categories, respectively, of all tracked cells from n = 5 datasets from the base medium condition imaged between ∼72-88hpa, i.e. around AP polarization. **E)** For two replicate datasets from base medium, individual tracks are displayed as vectors with the origin position set to 0,0. **F), G)** Pie charts as in **D)** for n = 4 datasets from the base+1i condition imaged and tracked between ∼72-83hpa **(F)** as well as for n = 3 datasets from the base+2i condition imaged and tracked between ∼96-110hpa **(G).**

**Fig. S4.**
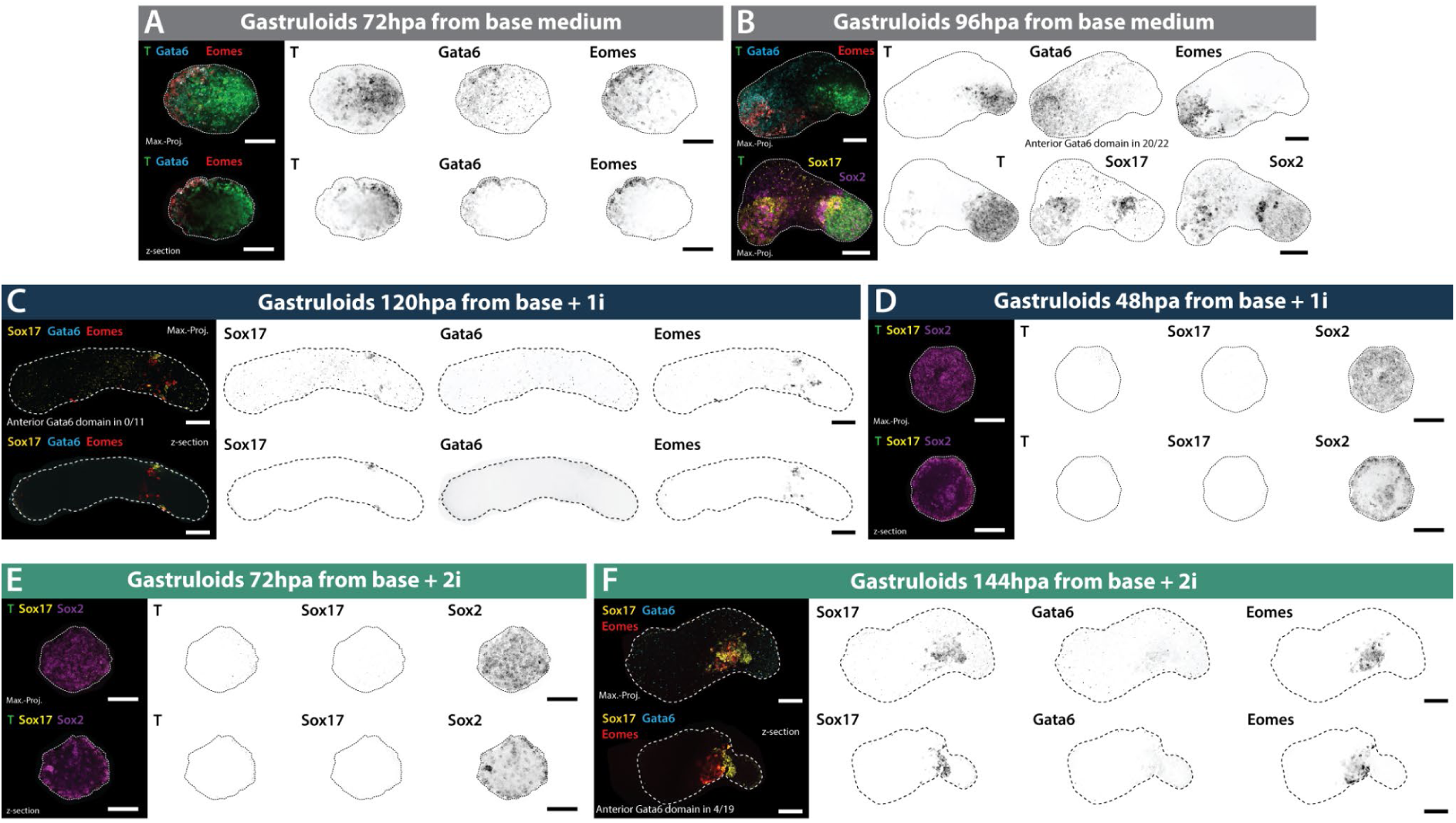
Tissue fate specification and localization differs between initial media conditions II. Representative images of HCR stainings (maximum intensity projections and single z-sections) of germ layer and axial patterning marker genes in gastruloids from three mESC media conditions (or pluripotent states) at comparable developmental timepoints. All scale bars are 100μm. **A), B)** Gastruloids from mESCs grown in base medium following T polarization **A)** and during or at maximum elongation **B)**. **C), D)** Aggregates from base+1i medium at maximum elongation **C)** and prior to CHIR99 application **D)**. **E), F)** Aggregates from base+2i before the CHIR99 pulse **E)** and at maximum elongation **F)**.

**Fig. S5.**
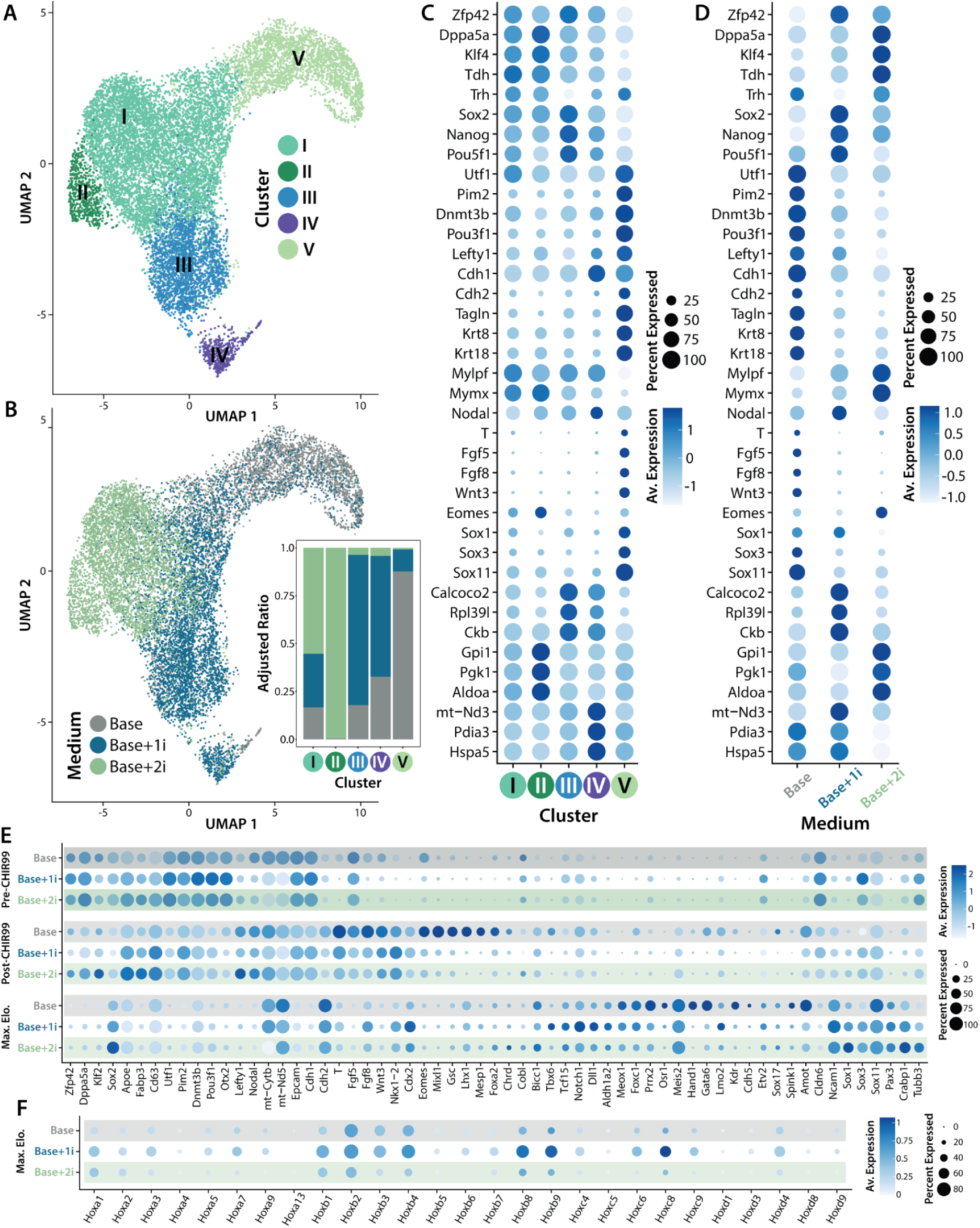
Single-cell transcriptomics delineate distinctions in lineage specification dynamics II. **A), B)** UMAP plots of the integrated mESC (0h) dataset. **A)** Color code is according to clustering analysis. **B)** Color code is according to the culture medium condition (i.e. pluripotent state). The inset on the right displays the adjusted ratios of cells of each medium condition in each of the 5 clusters. **C), D)** Dotplots displaying scaled average expression of various marker genes denoting cell state (naive pluripotent to early differentiating) and the percentage of cells expressing a given marker for either each cluster **C)** or medium condition **D)**. **E)** Dotplot of the integrated gastruloid dataset (see Figure 3) showing scaled average expression of cell state marker genes for all three source mESC media conditions and developmental timepoints. **F)** Dotplot of the maximum elongation timepoint dataset showing average expression of hox cluster genes for all three source mESC media conditions.

**Fig. S6.**
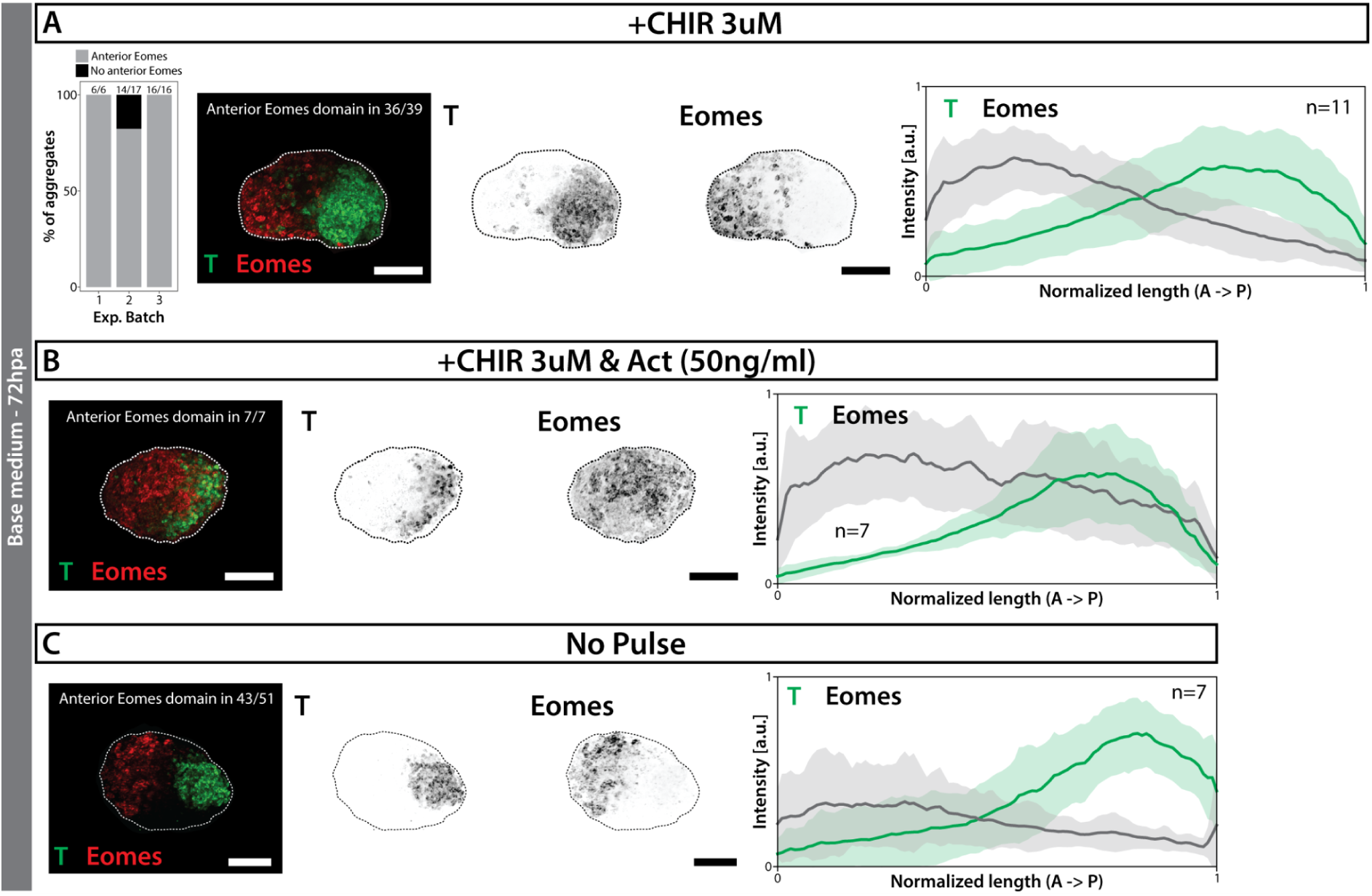
Aggregates from base medium exhibit highest inherent capacity to generate anterior mesoderm. Comparison of gastruloids from mESC cultured in base medium, subjected to a 24h pulse (between 48- 72hpa) of either CHIR99 (CHIR) as per the standard protocol **A)**, CHIR99 and ActivinA (Act) **B)**, or no pulse at all **C)**. The bar chart in **A)** (leftmost panel) depicts the percentage of aggregates from a given experimental batch exhibiting an anterior Eomes domain (also see Figure 5). The middle panel in **A)** or the left panels in **B), C)** show representative images (maximum intensity projections) of HCR stainings of T and Eomes with scale bars = 100μm. The rightmost panels display line plots of normalized fluorescence intensity of marker genes T and Eomes along the gastruloid’s AP axis (as demarcated by localized expression of T. Central lines denote the average across replicated, surrounding shades the standard deviation. Quantifications are based on sum intensity projections of the HCR data.

## Multimedia Files

**Movie S1** - Representative gastruloid from base medium. Sparse labeling was performed by mixing 5- 10% of 2KI reporter mESCs with WT mESCs (300 total cells). T signal via p2a-GFP is in green, the ubiquitous nuclear label via mKO2 in white. Numbers in top left denote the imaging time-window (hpa:min). Scale bar = 100μm.

**Movie S2** - Representative gastruloid from base+1i medium. Sparse labeling was performed by mixing 5-10% of 2KI reporter mESCs with WT mESCs (400 total cells). T signal via p2a-GFP is in green, the ubiquitous nuclear label via mKO2 in white. Numbers in top left denote the imaging time-window (hpa:min). Scale bar = 100μm.

**Movie S3** - Representative gastruloid from base+2i medium. Sparse labeling was performed by mixing 5-10% of 2KI reporter mESCs with WT mESCs (500 total cells). T signal via p2a-GFP is in green, the ubiquitous nuclear label via mKO2 in white. Numbers in top left denote the imaging time-window (hpa:min). Scale bar = 100μm.

